# YAP/TAZ are Crucial Regulator of Macrophage-mediated Pulmonary Inflammation and Fibrosis after Bleomycin-induced Injury

**DOI:** 10.1101/2023.05.24.542136

**Authors:** Masum M. Mia, Siti Aishah Binte Abdul Ghani, Dasan Mary Cibi, Hanumakumar Bogireddi, Wai Shiu Fred Wong, Manvendra K. Singh

**Author notes:** Corresponding author: Address for correspondence: Manvendra K. Singh; Program in Cardiovascular & Metabolic Disorders; Duke-NUS Medical School, 8 College Road, Office 08-15 Singapore 169857.; or.

## Abstract

Pulmonary fibrosis (PF) is the most common form of end stage interstitial devastating lung disease characterized by the scarring of lung due to excessive production of extracellular matrix (ECM). Recent studies have revealed the impact of macrophages in inflammation-induced fibrosis and distinct subsets of macrophages differentially contributes to the development of PF. However, the regulatory mechanisms and proinflammatory/profibrotc behaviour of heterogeneous population of lung macrophages during fibrogenesis remain incompletely understood. Here, we demonstrate the macrophage-specific role of Yes-associated protein (YAP) and transcriptional coactivator with PDZ-binding motif (TAZ) in the development of bleomycin-induced inflammation and PF in mice. Both YAP/TAZ are activated in lung macrophages of fibrotic patients and of mice after bleomycin-induced injury. Myeloid-specific genetic deletion of *Yap/Taz* resulted in reduced recruitment of monocyte-derived alveolar macrophages (Mo-AMs), leading to an impaired inflammatory response, reduced PF and improved regeneration of alveolar epithelial cells in bleomycin-injured lung. However, overexpression of *Yap* in macrophages augmented the Mo-AMs recruitment in lung leading to increased proinflammatory response, exacerbated fibrotic response and decreased regeneration of alveolar epithelial cells in bleomycin-injured lung. We demonstrate that YAP/TAZ regulate PF through the activation of macrophage recruitment driver C-C motif chemokine ligand 2 (CCL2) and blocking of CCL2 with neutralizing antibody prevented YAP-induced inflammatory and fibrotic response. We also demonstrate that YAP/TAZ regulate macrophage polarization as well as macrophage-fibroblasts crosstalk by regulating expression of Methyl-CpG–binding domain 2 (MBD2) during bleomycin-induced PF. Taken together, we show that YAP/TAZ are potent regulators of macrophage polarization, infiltration and macrophage–mediated proinflammatory/profibrotic response during PF.

## Introduction

Pulmonary fibrosis (PF) is an incurable interstitial lung disease with increasing mortality, characterized by the chronic inflammation, progressive scarring of lung due to excessive accumulation of extracellular matrix (ECM) between the alveoli’s or air sacs, and destruction of lung parenchyma. This results a decline in oxygen exchange function of lung, breathlessness and that eventually leads to organ failure. Number of factors are known to trigger PF including ageing, genetic disorders, autoimmune diseases, radiation, environmental factors and most recently the SARS-CoV-2 infection (Barratt, Creamer et al., 2018, Chen, Qiao et al., 2020, Lederer & Martinez, 2018). When the underlying cause of PF is unknown it termed as idiopathic pulmonary fibrosis (IPF), a progressive lung disease with complex pathophysiology primarily occurring in older adults and the median survival with IPF is 2-5 years after diagnosis (Ley, Collard et al., 2011). The availability of effective treatment including lung transplantation are limited to anti-fibrotic drug pirfenidone and nintedanib with significant impacts on patient survival and quality of life (Lederer & Martinez, 2018, Noble, Albera et al., 2011, Richeldi, du Bois et al., 2014); and there is an unmet clinical need for new therapeutic target. Despite the growing number of IPF patients, the pathological mechanism of the PF/IPF remains elusive.

Infiltration of inflammatory cells such as neutrophils, dendritic cells, leukocytes and macrophages into the injured site after organ injury including IPF are common. Whether inflammation contributes to IPF remains under debate, since anti-inflammatory agents are ineffective to improve clinical outcomes (Idiopathic Pulmonary Fibrosis Clinical Research, Raghu et al., 2012, Luppi, Cerri et al., 2004, Wuyts, Agostini et al., 2013). For example, nonselective depletion of both tissue-resident and monocyte-derived alveolar macrophages might worsen the fibrosis through further recruitment of monocytes that affect tissue homeostasis (Gibbings, Goyal et al., 2015, Janssen, Barthel et al., 2011). Therefore, understanding the contribution of specific cell populations and their interactions in the fibrotic niches are essential to prevent the pathogenesis of PF/IPF. Growing evidence indicates macrophages are crucial regulator of PF and distinct populations of lung macrophages differentially contribute to PF; however, the regulatory mechanisms and pro-inflammatory behaviour of distinct subpopulations underlying IPF pathology remain poorly understood.

In the lung, two distinct major populations of macrophages have been identified as alveolar macrophages (AMs) and interstitial macrophages (IMs). They are responsible to initiate innate immune response in the lung and involved to clearing the airways of pathogenic microbes, apoptotic cells, debris, and other pollutants through phagocytosis, (Hussell & Bell, 2014, Schyns, Bureau et al., 2018). After injury both AMs and IMs populations are expand in lung and studies suggest that circulating monocytes are principal source of newly derived macrophages, which play critical role in inflammation and different stages of fibrogeneis (Chakarov, Lim et al., 2019, Joshi, Watanabe et al., 2020, McCubbrey, Barthel et al., 2018, Misharin, Morales-Nebreda et al., 2017). Recent studies have evident that mouse AMs from bleomycin-induced PF are transcriptionally heterogeneous (McCubbrey et al., 2018). Similar transcriptional heterogeneity was described in AMs from IPF patients (Misharin et al., 2017), indicating the distinct AM subtypes may have a different role in the pathogenesis of PF. For example, targeted depletion of monocyte-derived alveolar macrophages (Mo-AMs) reduced Asbestos-induced lung fibrosis (Joshi et al., 2020). Indeed Misharin AV et al. (Misharin et al., 2017) showed that depletion of Mo-AMs through necroptosis during their differentiation reduced severity of bleomycin-induced PF; however depletion of tissue-resident AMs (TR-AMs) did not contribute to induce fibrosis. Similarly, by deleting the antiapoptotic protein c-Flip in AMs a genetic evidence revealed that Mo-AMs are causally related to severity of bleomycin-induced PF (McCubbrey et al., 2018). In contrast, a recent study evident that Mo-AMs actively participate into the resolution of bleomycin-induced lung fibrosis through ApoE/LRP1 mediated phagocytosis of collagen, suggesting Mo-AMs may have distinct functions in distinct stages of PF (Cui, Jiang et al., 2020). Recently, two distinct IMs lineage from monocyte derived tissue resident macrophages have been identified and depletion of Lyve1^high^MHCII^low^ IMs population exacerbated the bleomycin-induced PF (Chakarov et al., 2019), indicating the complexity of AMs and IMs functions in lung fibrogenesis and thus it is important to understand the role of mediators associated with the functional regulation of AMs and IMs in different phases of lung injury and repair.

Yes-associated protein (YAP) and transcriptional coactivator with PDZ-binding motif (TAZ) are the main transcriptional regulators of the Hippo signaling pathway known to regulate the development, homeostasis, and regeneration of numerous mammalian organs including lung (Fu, Plouffe et al., 2017). The existing evidence also suggests that they are crucial to modulate immune response by regulating both immune and non-immune cell (Guo, Zhao et al., 2017, Hagenbeek, Webster et al., 2018, Wang, Lu et al., 2016, Wang, Xie et al., 2017).Very recently we and others have demonstrated that YAP/TAZ are essential regulators of macrophage-polarization after organ injury (Mia, Cibi et al., 2020, Zhou, Li et al., 2019). For instance, macrophage-specific knockout of YAP relieves inflammatory bowel disease (Zhou et al., 2019). In the line, Macrophage specific deficiency of YAP/TAZ resulted in improved cardiac remodelling and function after myocardial infarction due to reduced cardiac fibrosis (Mia et al., 2020). Given these findings, we hypothesized that YAP/TAZ may regulate the phenotypic and functional behaviours of AMs and IMs at different phases of lung injury and thereby contributes to the pathogenesis of PF/IPF.

In this study, we have found that YAP and TAZ are activated in lung macrophages after bleomycin-induced injury and in fibrotic patients samples. Genetic deletion of *Yap/Taz* in macrophages impairs the expression of proinflammatory cytokines and chemokines in the injured mice lung. We also demonstrate an impaired production of profibrotic components and thereby reduced fibrosis in the lung due to *Yap/Taz* deletion. *Yap/Taz* inactivation further leads to a decrease in the number of Mo-AMs and IMs population in bleomycin-treated lung. The expression profiling reveals a reduced induction of proinflammatory and profibrotic factors by these Mo-AMs and IMs population. Moreover, our study demonstrate that *Yap/Taz*-deficient mice show improved alveolar epithelial regeneration after bleomycin-induced injury, evident by reduced loss of PDPN-expressing alveolar epithelial type-1 cells (AT1 cells) and surfactant protein C-expressing alveolar epithelial type-2 cells (AT2 cells) post-injury. Consistently, overexpression of YAP using constitutively active YAP mutant (*YAP^5SA^*) augmented the proinflammatory response in the lung by increasing the recruitments of Mo-AMs in the injured lung. Consequently, YAP overexpression aggravates lung fibrosis and alveolar epithelial cell damage which halted lung regeneration at post-bleomycin-injury. Mechanistically, our study revealed that YAP/TAZ regulate PF fibrosis through the activation of C-C motif chemokine ligand 2 (CCL2) that drives macrophage infiltration and blocking of CCL2 with neutralizing antibody abrogated YAP-induced fibrotic response at post bleomycin-injury. We also identified Methyl-CpG–binding domain 2 (MBD2) acts as downstream of YAP to induce myofibroblasts formation and ARG1+ profibrotic macrophage polarization during bleomycin-induced PF. Together, our findings demonstrate that YAP/TAZ modulate proinflammatory and regenerative response in the lung by regulating profibrotic macrophage behaviour.

## Results

### YAP/TAZ are activated in lung macrophages of pulmonary fibrotic patients and bleomycin-treated mice

To study whether YAP and TAZ are activated in human lung macrophages due to fibrosis, we first investigated the presence of macrophages in normal adult lung (considered as healthy human lung) and fibrotic lung from human and analysed the expression of YAP/TAZ in macrophages. Immunofluorescence analysis on healthy and fibrotic lung sections revealed a significant abundance of CD68-positive (CD68+) macrophages in fibrotic lung abundant with the expression of collagen type I (collagen I), compared healthy adult lung (Supplementary Figure 1). YAP and TAZ protein were detectable in the nucleus of CD68+ macrophages and compared to healthy lung there was a strong increase of YAP/TAZ in CD68+ macrophages on fibrotic patient lung (Figure 1A-B).

**Figure-1:**
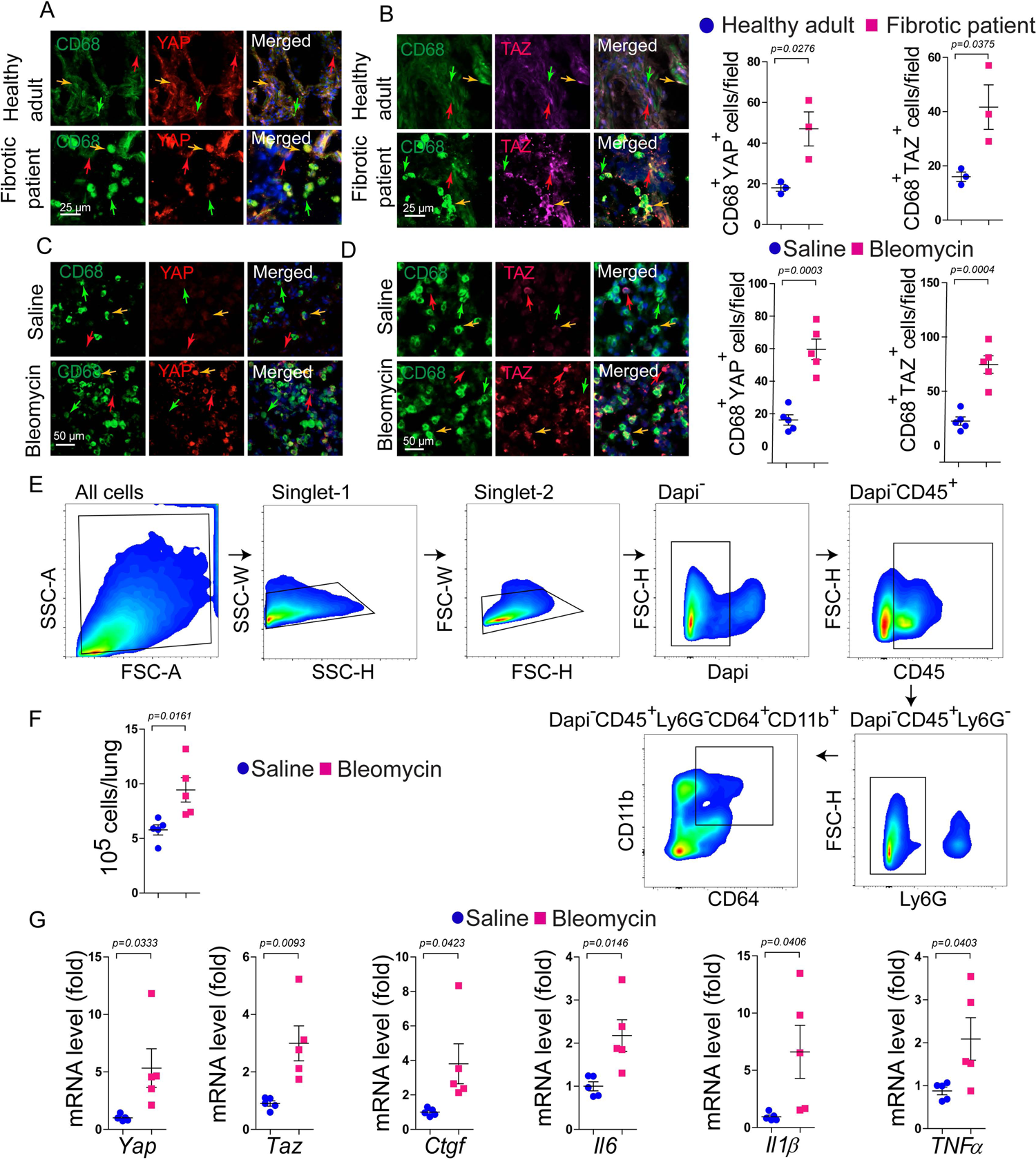
YAP/TAZ are activated in lung macrophages. (A-B) Immunofluorescence staining and quantification of YAP or TAZ co-stained with CD68 in lung sections from human patients diagnosed with pulmonary fibrosis compared with normal adult human (considered as healthy adult) lung (n = 3). (C-D) Immunofluorescence staining and quantification of YAP or TAZ co-stained with CD68 in wildtype mice lung sections at 7 days post-bleomycin injury compared to saline-treated sham controls (n = 5). (E) Gating strategy of flow cytometry analysis to sort the total lung macrophages (identified as Dapi^-^CD45^+^Ly6G^-^CD64^+^CD11b^+^ cells) using saline-treated sham controls (n = 5) and bleomycin-treated mice (n = 5) at 7 days. (D) mRNA analysis of *Yap*, *Taz*, *Ctgf*, *Il6*, *Il1β* and *Tnfα* on sorted total lung macrophages. The data are represented as the mean ± SEM; comparison by two-tailed unpaired t-test. *, *p* < 0.05; **, *p* < 0.01; ***, *p* < 0.001; NS, not significant.

To understand the macrophage-specific roles of YAP/TAZ in bleomycin-induced lung fibrosis, we examined their expression in mice at 7 days after intratracheal bleomycin administration and performed immunohistological analysis on lung sections. We observed enhanced nuclear presence of YAP and TAZ, since they act in the nucleus, in CD68+ macrophages in bleomycin-treated mice compared to saline-treated control mice (Figure 1C-D). Additionally, we observed increased expression of *Yap* and *Taz* mRNA level together with their target gene connective tissue growth factor (*Ctgf*) and pro-inflammatory macrophage marker interleukin-6 (*Il6),* interleukin-1beta *(Il1β)* and tumor necrosis factor alpha (*Tnfα*) in lung macrophages identified as Dapi^-^CD45^+^Ly6G^-^CD64^+^CD11b^+^ cells obtained through fluorescence-activated cell sorting (FACS) technique at 7 days after bleomycin instillation (Figure 1E-G). These preliminary observations suggesting that YAP/TAZ may contribute to lung macrophage function in response to injury.

### Inactivation of *Yap/Taz* impairs bleomycin-induced inflammation in the lung

To examine the role of YAP/TAZ in macrophage mediated inflammation and fibrosis in lung, we generated myeloid cell-specific *Yap/Taz* double knockout mice by crossing *Yap^flox/flox^;Taz^flox/flox^* mice with lysozyme-cre (*LysM^cre^*) mice that drive Cre recombinase activity in myeloid lineages, including macrophages. *Yap^flox/flox^;Taz^flox/flox^*and *LysM^cre^;Yap^flox/flox^;Taz^flox/flox^* mice are referred to as control and *Yap/Taz* double knockout (*dKO*), respectively. To investigate the role(s) of YAP/TAZ on bleomycin-induced inflammatory response, we treated the control and *dKO* mice for 7 and 14 days with bleomycin. To determine the effect of bleomycin, we also treated the control and *dKO* mice with saline and considered them as day 0 group. Immunohistological analysis on the bleomycin-injured lung sections showed reduced expression of YAP and TAZ in CD68+ cells of *LysM^cre^;Yap^flox/flox^;Taz^flox/flox^*mice suggesting an efficient deletion of YAP/TAZ in macrophages (Supplementary Figure 2). Subsequently, the bronchoalveolar lavage fluid (BALF) and perfused lungs were harvested for further analysis. Then we observed whether the presence of macrophage was affected after injury due to deletion of *Yap/Taz*. There was no change in CD68+ macrophage numbers in saline treated control *versus dKO* lung. Certainly, bleomycin-induced injury augmented the presence of CD68+ macrophages in the control lung at 7 and 14 days respectively, however *Yap/Taz* deficiency halted the macrophage infiltration (Figure 2A-B). Next, we measured the expression of pro-inflammatory cytokines such as IL1β, IL6 and TNF-α in BALF using enzyme-linked immunosorbent assay (ELISA). Compared to saline treatment, bleomycin-mediated injury led to the excessive production of IL1β, IL6 and TNF-α in BALF from control animals at day 7 and day 14, however *Yap/Taz*-deficient mice abrogated the production of these cytokines compared to control mice (Figure 2C). To understand the function of YAP/TAZ on inflammatory reaction further we examined the mRNA expression of pro-inflammatory cytokines and chemokines in lung tissue. Indeed, quantitative reverse-transcription PCR (qRT-PCR) analysis using lung tissue revealed that pro-inflammatory cytokines and chemokines such as *Il6*, *Il1β*, nitric oxide synthase-2 (*Nos2*), hypoxia-inducible factor 1-alpha (*Hif1α*), C-C motif chemokine ligand 2 (*Ccl2*), C-C motif chemokine receptor 2 (*Ccr2*), C-X3-C Motif chemokine ligand 1 (*Cx3cl1*), C-X3-C Motif chemokine Receptor 1 (*Cx3cr1*), and C-C motif chemokine ligand 17 (*Ccl17*) were downregulated in *dKO*-lung compared to controls at 14 days after bleomycin instillation. In addition, the expression of profibrotic macrophage genes such as Arginase 1 (*Arg1*), Resistin-like molecule alpha or found in inflammatory zone protein 1 (*Fizz1*) and YAP/TAZ target gene *Ctgf* were also dampened in *dKO*-lung compared to controls (Figure 2). No difference was detected between control and *dKO* lungs for these genes after saline instillation (Figure 2D).

**Figure 2:**
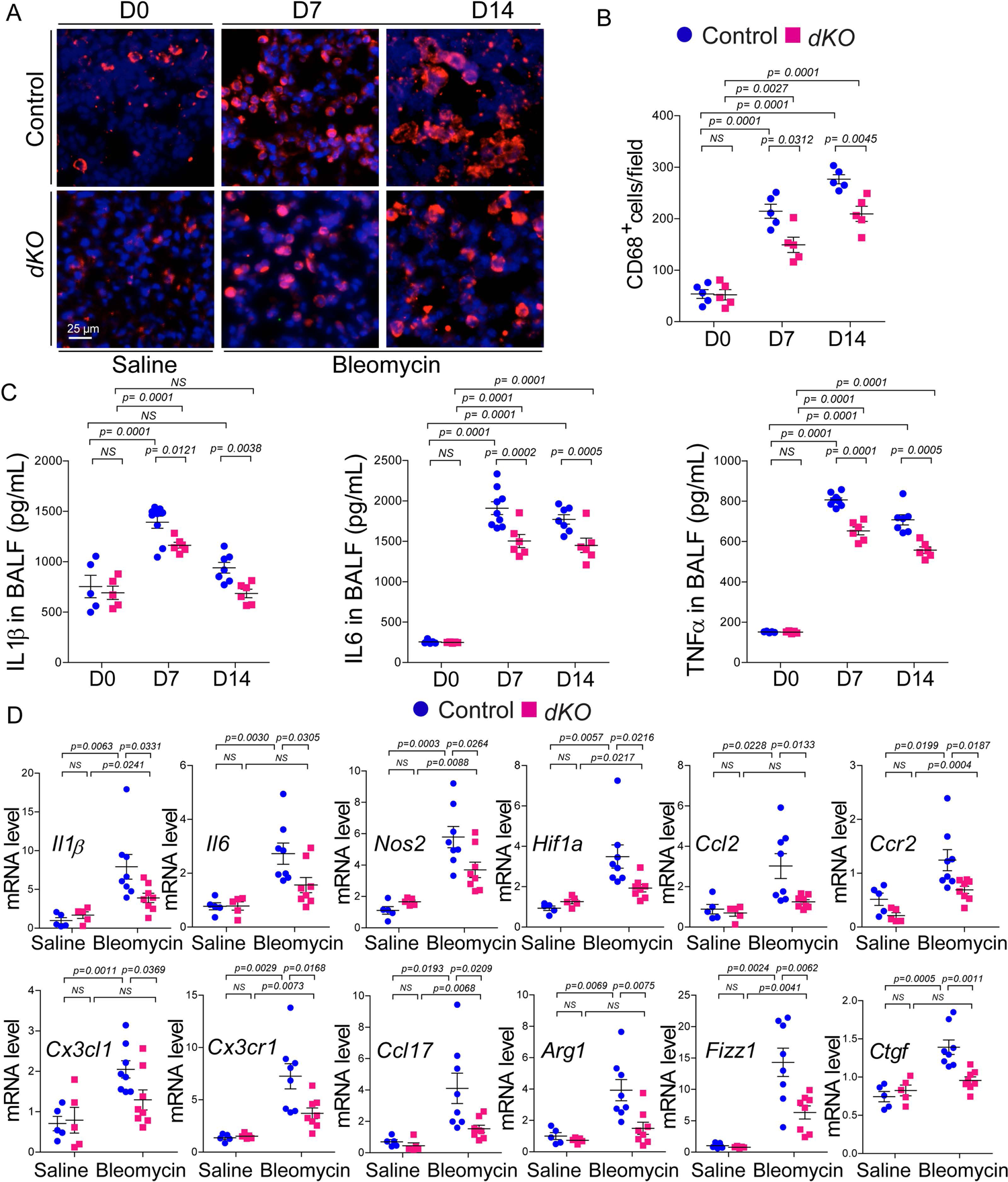
Macrophage-specific *Yap/Taz* inactivation impairs bleomycin-induced inflammation in mice lung. (A-B) Immunostaining and quantification of CD68 positive macrophage infiltration on the lung treated with saline considered as day 0 (D0) or bleomycin for day 7 (D7) and day 14 (D14) using control and *dKO* mice (n = 5). (C) Enzyme-linked immunosorbent assay (ELISA) to measure the protein release of pro-inflammatory cytokines IL1β, IL6 and TNF-α in bronchoalveolar lavage fluid (BALF) collected from control and *dKO* mice (n = 5-6) treated with saline (D0) or bleomycin for 7 and 14 days respectively. (D) Real-time qPCR for pro-inflammatory cytokines and chemokines such as *Il6*, *Il1β*, *Nos2, Hif1α, Ccl2, Ccr2, Cx3cl1, Cx3cr1*, *Ccl17, Arg1* and *Fizz1*; with YAP/TAZ target gene *Ctgf* using lung tissue RNA isolated from control and *dKO* mice (n = 5-8) treated with saline or bleomycin for 14 days. The data are represented as the mean ± SEM; comparison by two-tailed unpaired t-test. *, *p* < 0.05; **, *p* < 0.01; ***, *p* < 0.001; NS, not significant.

Growing evidence indicates macrophages are crucial regulator of bleomycin-induced lung inflammation, therefore we hypothesised that YAP/TAZ may regulate the pro-inflammatory behaviour of lung macrophages. In the lung, two major populations of macrophages have been identified as AMs and IMs. Given that both AMs and IMs populations are increasing after injury in lung and evident suggest that circulating monocytes are major origin of newly infiltrating macrophages to contribute into AMs and IMs phenotype, which play crucial role in inflammation and inflammation-mediated fibrogeneis (Chakarov et al., 2019, Joshi et al., 2020, McCubbrey et al., 2018, Misharin et al., 2017). We therefore examined whether deficiency of *Yap/Taz* affects the recruitments and functions of monocyte-derived macrophages after bleomycin administration in mice. Flow cytometry analyses on enzymatically digested lung tissue demonstrated a strong increase of Mo-AMs identified as CD45^+^Ly6G^-^CD64^+^CD11b^low^Siglec-F^low^ in control lung after bleomycin-induced injury at day 7 and day 14 respectively, however the number of Mo-AMs population were significantly reduced in *dKO* lung compared to controls (Figure 3A-B). Interestingly, the number of TR-AMs identified as CD45^+^Ly6G^-^CD64^+^CD11b^low^Siglec-F^high^ were decreased at day 7 for both control and *dKO* group due to bleomycin treatment; however at day 14, this population was unaffected after bleomycin-induced injury and no alteration was observed in response to *Yap/Taz* deletion (Figure 3C). The number of IMs identified as CD45^+^Ly6G^-^CD64^+^CD11b^high^Siglec-F^-^ showed a time-dependent alteration after bleomycin-injury. At day 7 after injury, the number of IMs population were significantly reduced both in control and *dKO* lung, however no differences were noticed due to loss of *Yap/Taz*. At day 14 after bleomycin administration, there was a modest increase of IMs population in control lung, however the number of this population were less in *dKO* lung compared to controls (Figure 3D). Together, myeloid-specific *Yap/Taz* inactivation leads to a decrease in Mo-AMs and IMs population in bleomycin-treated mice.

**Figure 3:**
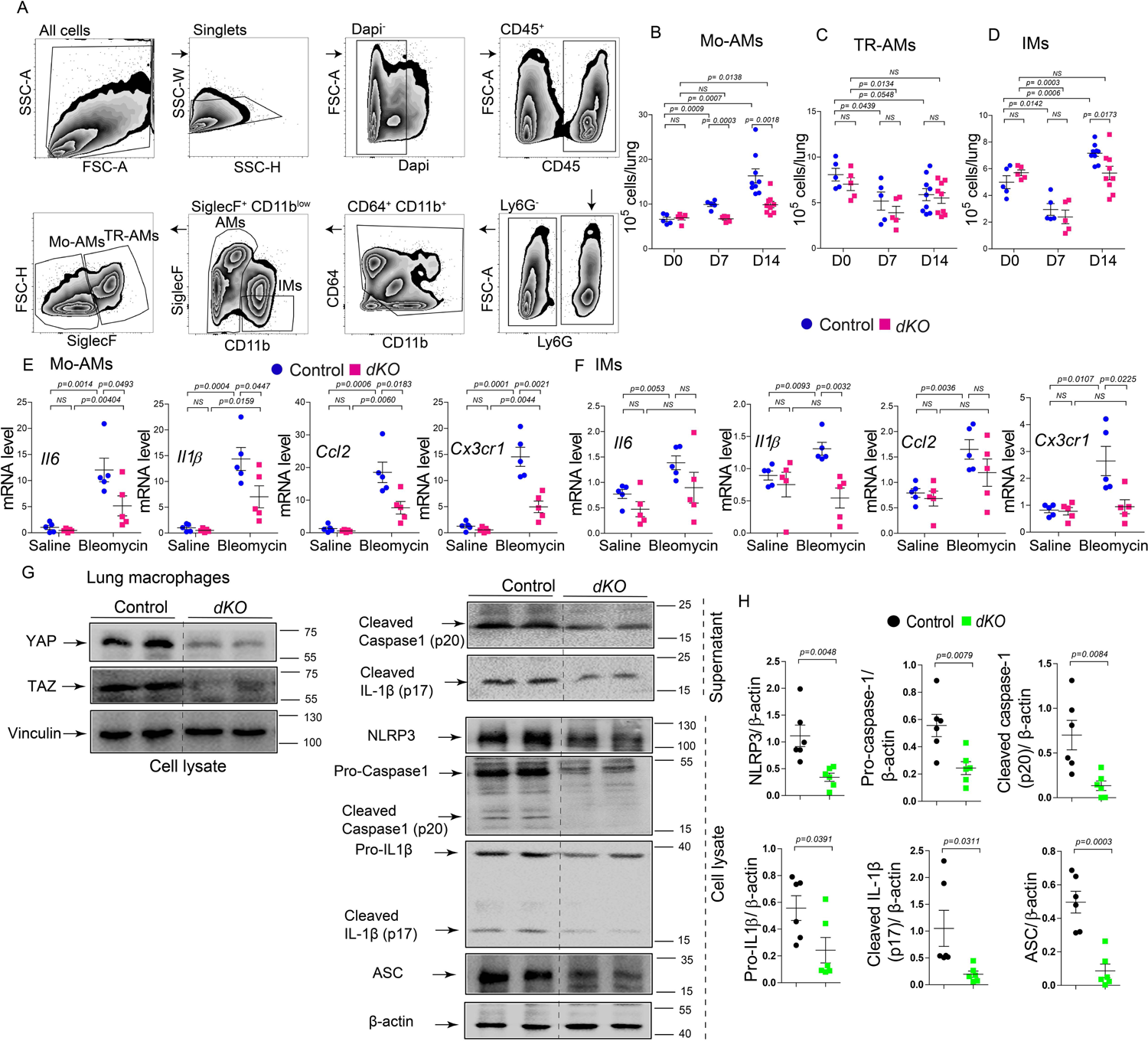
Macrophage-specific *Yap/Taz* inactivation leads to decrease in Mo-AMs and IMs population after bleomycin-induced injury. (A-D) Gating strategy and flow cytometry analysis to sort Mo-AMs (identified as CD45^+^Ly6G^-^ CD64^+^CD11b^low^Siglec-F^low^), TR-AMs (identified as CD45^+^Ly6G^-^ CD64^+^CD11b^low^Siglec-F^high^) and IMs (CD45^+^Ly6G^-^CD64^+^CD11b^high^Siglec-F^-^) using lung from control and *dKO* mice (n = 5-9) treated with saline (D0) or bleomycin for 7 (D7) and 14 days (D14). (E-F) Real-time qPCR for pro-inflammatory cytokines and chemokines such as *Il6*, *Il1β*, *Ccl2*, and *Cx3cr1* on sorted Mo-AMs or IMs from control and *dKO* mice (n = 5) treated with saline or bleomycin for 14 days. (G-H) Immunoblot analysis of YAP, TAZ, cleaved caspase-1, cleaved IL-1β, NLRP3, pro-caspase-1, pro-IL-1β and ASC using cell lysates or supernatants as indicated from cultured lung macrophages isolated through flow cytometry (identified as Dapi^-^ CD45^+^Ly6G^-^CD64^+^CD11b^+^ cells) on bleomycin treated control and *dKO* mice (n = 5-6). (H) Quantification of indicated proteins relative to β-actin. The data are represented as the mean ± SEM; comparison by two-tailed unpaired t-test. *, *p* < 0.05; **, *p* < 0.01; ***, *p* < 0.001; NS, not significant.

Since lack of Yap/Taz did not affect the numbers of TR-AMs after bleomycin injury, we next characterized the gene expression patterns of proinfllmmatory/profibrotic cytokines and chemokines on fluorescence-activated cell sorting (FACS)-sorted Mo-AMs and IMs (Figure 3E-F). The expression profiling displayed an upregulation of *Il6 (*∼ 10-fold), *Il1β* (∼ 15-fold), *Ccl2* (∼ 18-fold), and *Cx3cr1 (*∼ 14-fold) in Mo-AMs after bleomycin treatment in control lung compared to saline-treated lung, however *Yap/Taz* deficiency prevented the elevation of these proinfllmmatory/profibrotic factors after bleomycin-induced injury. No difference were found in saline treated control versus *dKO* M0-AMs (Figure 3E). In IMs, the mRNA level of *Il6 (*∼ 1.5-fold), *Il1β* (∼ 1.4-fold), *Ccl2* (∼ 1.6-fold), and *Cx3cr1 (*∼ 2.5-fold) were significantly elevated after bleomycin injury in control lung; however fold-induction of these cytokine/chemokines were less compared to Mo-AMs. Similar to Mo-AMs, IMs from *dKO* lung revealed a reduced expression *Il6*, *Il1β,* and *Cx3cr1*; however the *Ccl2* expression was unaffected. In saline treated group, *Yap/Taz* deficient IMs showed no alteration for the expression of *Il6*, *Il1β*, *Ccl2,* and *Cx3cr1* compared to controls (Figure 3F). These findings suggested that YAP/TAZ has a novel role in regulating lung macrophage heterogeneity during injury.

To study the macrophage-mediated lung inflammation further, we have evaluated the role of YAP/TAZ in inflammasome activation. Activation of NLRP3 (nucleotide-binding oligomerization domain-like receptor containing pyrin domain 3) inflammasome in alveolar macrophages significantly contributes to induce the lung inflammation and injury through the release of inflammatory mediator IL-1β via caspase-1 (Wu, Yan et al., 2013). To test the role of YAP/TAZ in NLRP3 activation, we have cultured the sorted the lung macrophages identified as Dapi^-^CD45^+^Ly6G^-^ CD64^+^CD11b^+^ cells from control and *dKO* mice treated with bleomycin for 14 days and measured the expression of inflammasome factors NLRP3, caspase-1, IL-1β and ASC. Immunoblot analysis revealed that deficiency of YAP and TAZ in lung macrophages markedly decreased the released of cleaved caspase-1 and cleaved IL-1β in cultured supernatant. Similarly in cell lysate, the protein expression of NLRP3, pro-caspase-1, cleaved caspase-1, pro-IL-1β, cleaved IL-1β and ASC were significantly suppressed due to *Yap/Taz* deletion (Figure 3G-H). NLRP3 is essential for caspase-1-dependent maturation of IL-1β during inflammasome activation (Kayagaki, Warming et al., 2011). Based on our findings, YAP/TAZ deficiency decreased the release of caspase-1 and IL-1β by lung macrophages, suggesting that loss of YAP/TAZ impairs the NLRP3 inflammasome activation.

### Inactivation of *Yap/Taz* attenuates bleomycin-induced lung fibrosis

In order to understand the YAP/TAZ dependent functions of macrophages in lung fibrosis, we performed histochemical, immunohistological, immunoblot and qRT-PCR analyses on lung sections in bleomycin-treated control *versus dKO* mice. Bleomycin-induced injury led a development of lung fibrosis during a 7 and 14 day time course experiment (Supplementary Figure 3) (Moore & Hogaboam, 2008) and deletion of YAP/TAZ attenuated fibrosis in *dKO* lung compared to littermate controls as indicated with reduced Sirius red positivity at day 7 and day 14 post-injury analyzed with Ashcroft score (Figure 4A-B). Next, we measured the profibrotic collagen content in lung tissue as determined by hydroxyproline assay. Bleomycin-induced injury augmented the hydroxyproline content in control lung, however there was a decrease of hydroxyproline content in *dKO* lung compared to control and that almost reached baseline expression at 14 days post-injury (Figure 4C). Consistent with these findings, we have examined the expression of most abundant fibrogenic markers Collagen type I (collagen I) and myofibroblasts marker aSMA on lung tissue. Subsequently, we observed a profound expression of collagen I and aSMA in control lungs after bleomycin administration compared to saline treatment; however collagen I and aSMA level were reduced in *dKO* lung as detected through immunofluorescence and immunoblot analysis at day 14 post-injury. No significant changes were observed between saline-treated control and *dKO* lung (Supplementary Figure 4A-C). Similarly, gene expression analyses revealed pro-fibrotic ECM related genes such as collagen type I, III, and fibronectin 1 (*Col1a1*, *Col3a1* and *Fn1*), and myofibroblast-related genes such as smooth muscle actin alpha 2 (*Acta2*) were downregulated in *dKO* lung compared to controls at 14 days after bleomycin instillation (Figure 4D). Compared to saline, bleomycin treatment notably increased the expression of these genes, although no difference was detected within saline treated control *versus dKO* lung (Figure 4D). These findings strongly suggest that YAP/TAZ are enriched in the lung macrophages during bleomycin-induced fibrosis and act as crucial regulator of macrophage heterogeneity and macrophage functions that promote lung fibrosis.

**Figure 4:**
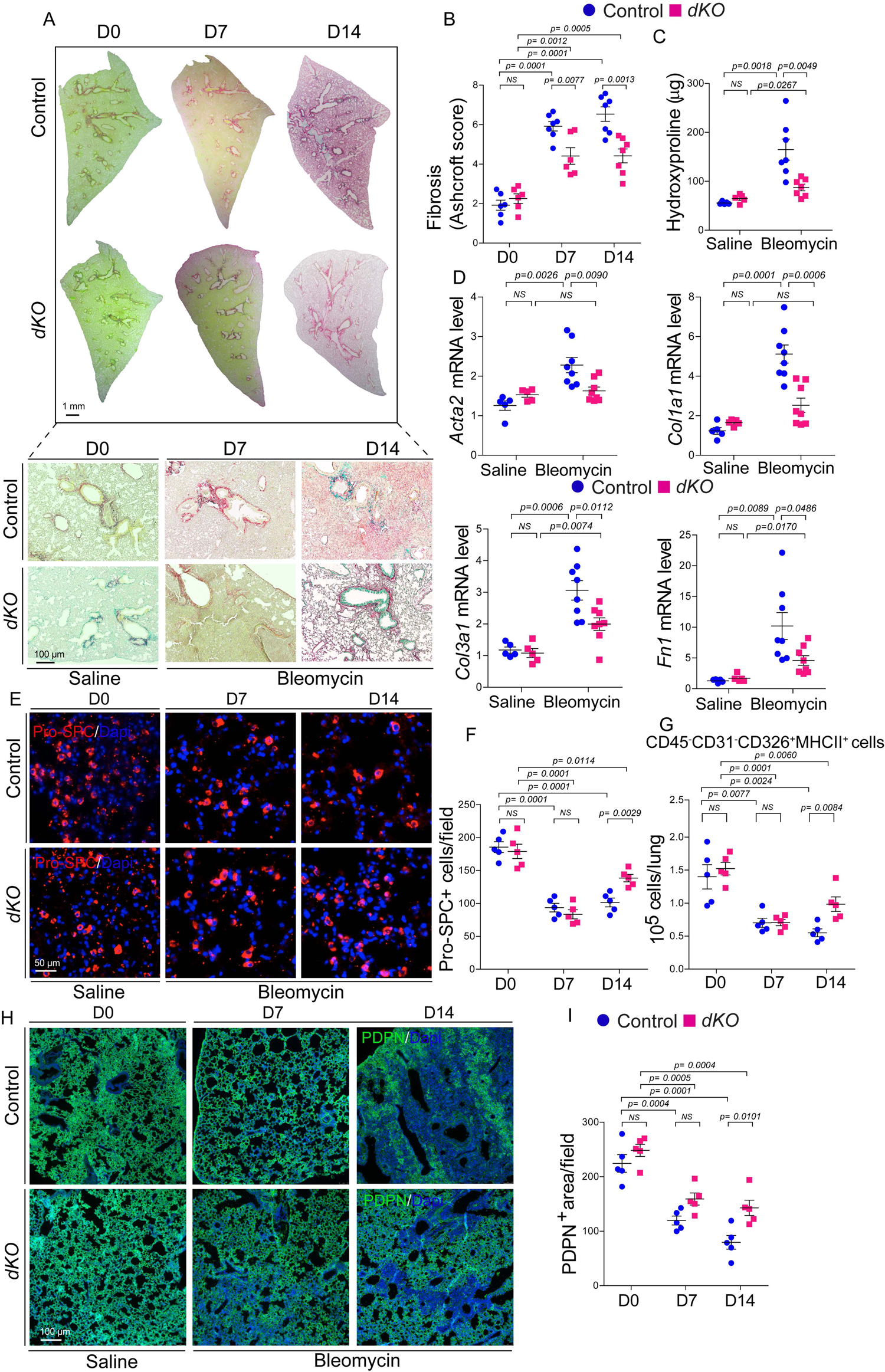
Macrophage-specific *Yap/Taz* inactivation reduces lung fibrosis after bleomycin-induced injury. (A-B) Sirius Red/Fast Green staining and quantification (presented as Ashcroft score) on lung sections from control and *dKO* mice (n = 6-7) treated with saline (D0) or bleomycin for 7 and 14 days respectively. (C) Quantification of hydroxyproline content using lung tissue from control and *dKO* mice (n = 5-7) treated with saline or bleomycin for 14 days. (D) Real-time qPCR for profibrotic genes *Acta2, Col1a1, Col3a1* and *Fn1* on lung tissue from control and *dKO* mice (n = 5-8) treated with saline or bleomycin for 14 days. (E-F) Immunostaining and quantification of Pro-SPC^+^ AT2-cells in control *versus dKO* mice lung (n = 5) treated with saline (D0) or bleomycin for 7 (D7) and 14 days (D14). (G) Flow cytometry analysis to determine the changes in the number of AT2-cells identified as CD45^−^CD31^−^CD326^+^MHCII^+^ in *dKO* lungs compared to controls (n = 5) after saline or bleomycin treatment for 7 and 14 days. (H-I) Immunostaining and quantification of PDPN^+^ AT1-cells in *dKO* lungs compared to controls (n = 5) after saline or bleomycin treatment for 7 and 14 days. The data are represented as the mean ± SEM; comparison by two-tailed unpaired t-test. *, *p* < 0.05; **, *p* < 0.01; ***, *p* < 0.001; NS, not significant.

### Inactivation of *Yap/Taz* protects bleomycin-induced alveolar epithelial cell damage

In the lung alveoli, alveolar epithelial type-2 cells (AT2 cells) serve as resident stem cells that can proliferate and differentiate into alveolar epithelial type-1 cells (AT1 cells) which are essential for gas exchange and to regenerate new alveoli after lung injury (Adamson & Bowden, 1974, Barkauskas, Cronce et al., 2013). Dysfunction or damage of alveolar AT2 cells has been reported to drives the progression of pulmonary fibrosis (Selman & Pardo, 2020, Sisson, Mendez et al., 2010). Therefore we aim to examine the regenerative potential of alveolar epithelial cells due to inactivation of *Yap/Taz* in macrophages of mouse lung exhibiting bleomycin-induced pulmonary fibrosis. To understand whether YAP/TAZ contributes to alveolar epithelial repair or prevent epithelial damage *in vivo*, *dKO* mice in parallel with their littermate controls were exposed to bleomycin-induced lung injury for 7 and 14 days respectively. Lung tissues were harvested to perform the immunohistochemistry, flow cytometry, and qRT-PCR analysis. We observed a strong damage in epithelial cells after bleomycin administration into the lung as indicated by the significant loss of AT1 and AT2 cells after day 7 and 14 post-injury compared saline treated lung (Figure 4E-I). Immunofluorescence analysis revealed that the number of surfactant protein C-expressing AT2 cells, as indicated by the pro-SPC antibody immunostaining, were strongly reduced at day 7 and day 14 post-injury in both control and *dKO* lung. No alterations in AT2 cell loss were detected between controls *versus dKO* group after day 7 of injury; In contrast, there was an increase of pro-SPC+ AT2-cells in *dKO* lung compared to controls at day 14 post injury (Figure 4E-F). Consistent with this finding, flow cytometry analysis showed a similar results; a strong decrease in the number of AT2 cells as identified by CD45^−^CD31^−^CD326^+^MHCII^+^ cells in both control and *dKO* bleomycin-injured lung at day 7 and 14 respectively. However, at 14 days post-injury there was a significant increase in the number of CD45^−^CD31^−^CD326^+^MHCII^+^ cell population in *dKO* lung compared with that control lungs (Figure 4G) (Supplementary Figure 5A). This increase in AT2 cells in *dKO* lung was correlated the mRNA levels of *Sftpc*, an AT2 cell marker, at day 14 post-injury (Supplementary Figure 5B). We next quantified the damage of AT1 cells and subsequently observed that compared to saline treatment there was an impaired AT1 cell numbers due to bleomycin injury as detected by the immunofluorescence staining of PDPN-positive (AT1 cell marker) area. We found an increase PDPN^+^ area in *dKO* lung compared with that control lungs at 14 days post-injury, however no difference was found between control and *dKO* lung at day 7 after injury (Figure 5H-I). Similarly, expression of another AT1 marker *Cav1* was downregulated in control and *dKO* lung after bleomycin treatment compared to their saline treated lungs, although the decrease of *Cav1* mRNA level in *dKO* was significantly prevented compared controls after injury at day 14. Saline treatment did not affect the expression of *Cav1* between these groups (Supplementary figure 5C). Collectively, these findings suggest that mice that lacked with *Yap/Taz* in macrophages showed improved epithelial regenerative response in the lung after bleomycin-induced injury.

**Figure 5:**
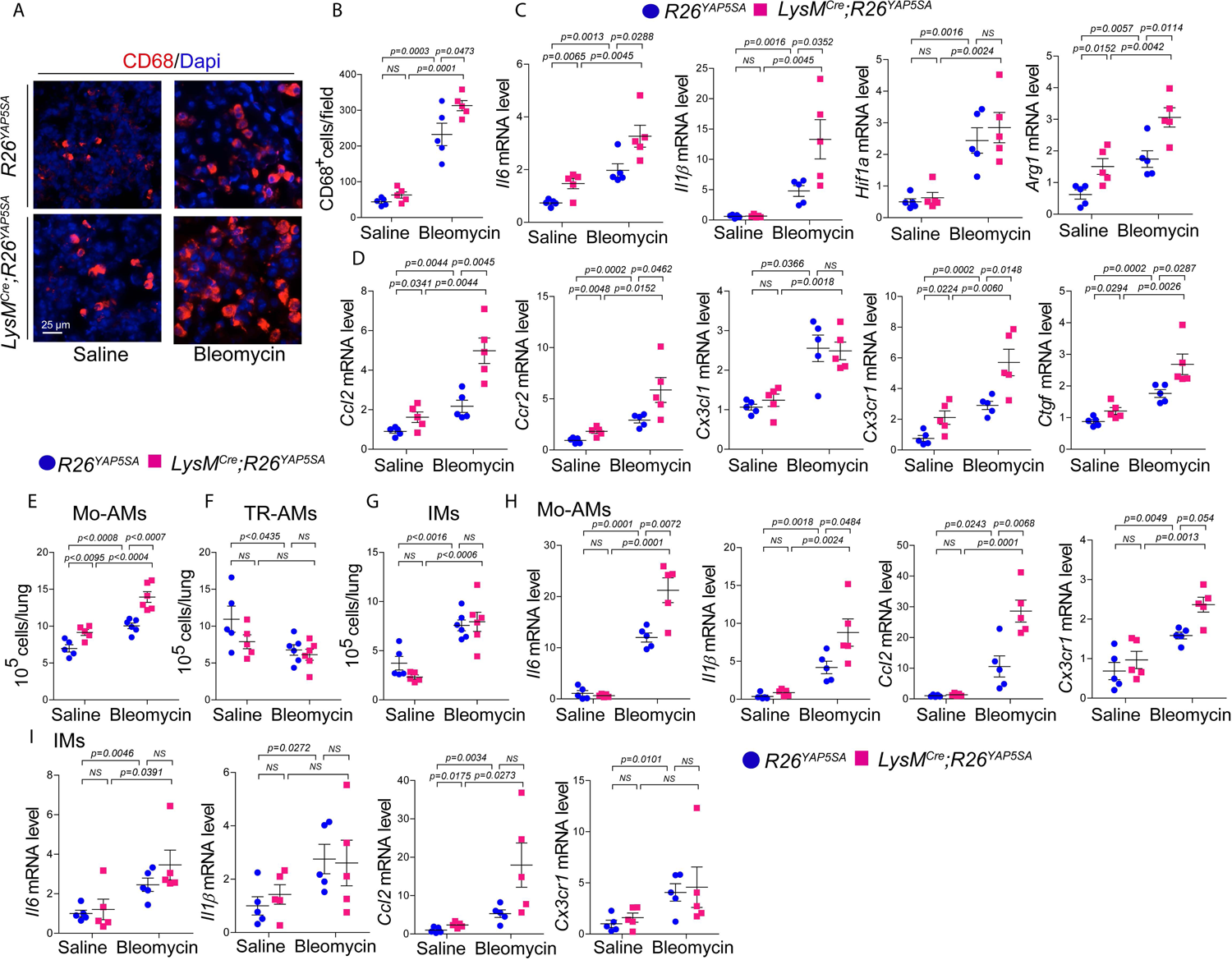
Macrophage-specific *Yap* overexpression augments bleomycin-induced inflammatory response in mice lung. (A-B) Immunostaining and quantification of CD68^+^ macrophage infiltration on the lung from *R26^YAP5SA^* and *LysM^Cre^;R26^YAP5SA^*mice (n = 5) treated with saline or bleomycin for 14 days. (C-D) Real-time qPCR for pro-inflammatory cytokines and chemokines such as *Il6*, *Il1β*, *Hif1α, Arg1, Ccl2, Ccr2, Cx3cl1, and Cx3cr1*; with hippo target gene *Ctgf* using lung tissue RNA isolated from *R26^YAP5SA^* and *LysM^Cre^;R26^YAP5SA^*mice (n = 5) treated with saline or bleomycin for 14 days. (E-G) Flow cytometry analysis to sort the Mo-AMs (identified as CD45^+^Ly6G^-^CD64^+^CD11b^low^Siglec-F^low^), TR-AMs (identified as CD45^+^Ly6G^-^CD64^+^CD11b^low^Siglec-F^high^) and IMs (CD45^+^Ly6G^-^ CD64^+^CD11b^high^Siglec-F^-^) using lung from *R26^YAP5SA^* and *LysM^Cre^;R26^YAP5SA^* mice (n = 5-6) treated with saline or bleomycin for 14 days. (H-I) Real-time qPCR for pro-inflammatory cytokines and chemokines such as *Il6*, *Il1β*, *Cx3cr1*, and *Ccl2* on sorted Mo-AMs or IMs from *R26^YAP5SA^* and *LysM^Cre^;R26^YAP5SA^* mice (n = 5) treated with saline or bleomycin for 14 days. The data are represented as the mean ± SEM; comparison by two-tailed unpaired t-test. *, *p* < 0.05; **, *p* < 0.01; ***, *p* < 0.001; NS, not significant.

### Macrophage-specific YAP activation augments bleomycin-induced inflammatory response and fibrosis in lung

To further investigate the role of YAP in inflammation induced fibrosis, we activated YAP in myeloid cells using a conditional Rosa26 allele, *R26^YAP5SA^*, and *LysM^Cre^* mice. This allele (*LysM^Cre^;R26^YAP5SA^*) enabled us to show the *in vivo* expression of a constitutively active form of YAP in macrophages. A strong nuclear presence of YAP in CD68-positive macrophages of *LysM^Cre^;R26^YAP5SA^* lung was observed, suggesting YAP activation in lung macrophages (Supplementary figure 6). To determine the effect of YAP on macrophage-mediated inflammation and fibrosis, we have treated the mice with saline or bleomycin for 14 days and lung were analysed with qRT-PCR, flow cytometry, immunoblot and immunostaining. We investigated whether the presence of macrophage was affected after bleomycin-induced injury due to overexpression of YAP. No changed was observed in CD68+ macrophage numbers in saline treated *R26^YAP5SA^* and *LysM^Cre^;R26^YAP5SA^* lung. As expected, bleomycin-induced injury augmented the presence of CD68+ macrophages in the *R26^YAP5SA^* lung at 14 days after injury and interestingly the number of CD68+ macrophages increased further in *LysM^Cre^;R26^YAP5SA^* lung, suggesting YAP promotes the macrophage infiltration after injury (Figure 5A-B). Next, we defined the expression of proinflammatory signatures. qRT-PCR analysis revealed that the expression of pro-inflammatory macrophage cytokines such as *Il6 and Il1β*; chemokines such as *Ccl2, Ccr1* and *Cx3cl1 Cx3cr1*; profibrotic macrophage gene *Arg1*, hypoxia regulator *Hif1α* and hippo target gene *Ctgf* were affected in lung due to YAP overexpression (Figure 5C-D). In saline treated mice, we observed elevated level of *Il6, Arg1, Ccl2, Ccr2, Cx3cr1* and *Ctgf* in *LysM^Cre^;R26^YAP5SA^* lung compared to *R26^YAP5SA^* lung; however no difference were detected *for Il1β, Cx3cl1*, and *Hif1α*. Bleomycin administration induced the gene levels of *Il6*, *Il1β*, *Arg1, Ccl2, Ccr2, Cx3cr1*, and *Ctgf* in *R26^YAP5SA^* lung and their expression were further augmented in YAP overexpressing *LysM^Cre^;R26^YAP5SA^*lung. Similarly the expression of *Hif1α* and *Cx3cl1* were enhanced after bleomycin-induced injury in *R26^YAP5SA^* mice, interestingly their expression were not altered due to YAP overexpression at tissue level (Figure 5C-D). To understand whether YAP overexpression is associated with the infiltration of macrophages, we have sorted the lung macrophages with a similar strategy used for *dKO* model above (Figure 3A). Flow cytometry analysis showed that the numbers of Mo-AMs population were significantly increased in *LysM^Cre^;R26^YAP5SA^*lung compared to *R26^YAP5SA^* lung, while the number of TR-AMs and IMs were unchanged between this groups (Figure 5E-G). To determine the molecular changes associated with YAP activation, we have evaluated the mRNA levels of proinfllmmatory/profibrotic cytokines and chemokines on sorted Mo-AMs and IMs. We observed an enhanced expression levels of *Il6*, *Il1β*, *Ccl2*, and *Cx3cr1* in Mo-AMs derived from *LysM^Cre^;R26^YAP5SA^* lung compared *R26^YAP5SA^* lung after bloemycin injury; although no alteration was noticed in saline treated group (Figure 5H). Similarly in IMs, bleomycin induced the gene levels of *Il6*, *Il1β*, *Ccl2*, and *Cx3cr1* in *R26^YAP5SA^* lung; however compared to *R26^YAP5SA^* lung, *LysM^Cre^;R26^YAP5SA^*lung did not reveal any alteration for the expression of these genes (Figure 5I). In saline treated group, the expression of *Ccl2* was evaluated in *LysM^Cre^;R26^YAP5SA^* lung compared to *R26^YAP5SA^* lung, however no change was detected for *Il6*, *Il1β*, and *Cx3cr1*.

Next, we studied whether macrophage-specific activation of YAP has any effect on lung fibrosis at baseline and post-injury. Sirius red staining showed an increase in fibrosis presented as Ashcroft score due to activation of YAP in *LysM^Cre^;R26^YAP5SA^* lung both at baseline and 14 post-injury with bleomycin (Figure 6A-C). Consistent with the Ashcroft fibrotic score, hydroxyproline assay revealed an augment of collagen content in *LysM^Cre^;R26^YAP5SA^*lung compared to *R26^YAP5SA^* lung both at baseline and post-injury condition (Figure 6D). Next, we observed whether pro-fibrotic genes were affected in the lung tissue due to YAP overexpression. Indeed, overexpression of YAP enhanced the myofibroblast marker *Acta2* and ECM genes *Col1a1* and *Col3a1* in bleomycin treated *LysM^Cre^;R26^YAP5SA^*mice compared to *R26^YAP5SA^* mice. We also detected an upregulation of *Col1a1* and *Col3a1* in saline treated *LysM^Cre^;R26^YAP5SA^*lung compared to control, although no alteration was detected for *Acta2* expression in these groups (Figure 6E). Immunoblot analysis revealed the similar observation, bleomycin treatment induced the protein expression of collagen I and aSMA in *LysM^Cre^;R26^YAP5SA^* lung compared to *R26^YAP5SA^* lung, suggesting the YAP mediated upregulation of these proteins post-injury. In saline treated group, we observed an elevated levels of collagen I expression in *LysM^Cre^;R26^YAP5SA^*lung compared to *R26^YAP5SA^* lung, although no expressional difference was found for aSMA between these groups (Figure 6F-H). Collectively, these findings suggest that overexpression of YAP in the lung macrophages during bleomycin-induced injury may regulates of macrophage recruitments, heterogeneity, and functions associated with lung fibrosis.

**Figure 6:**
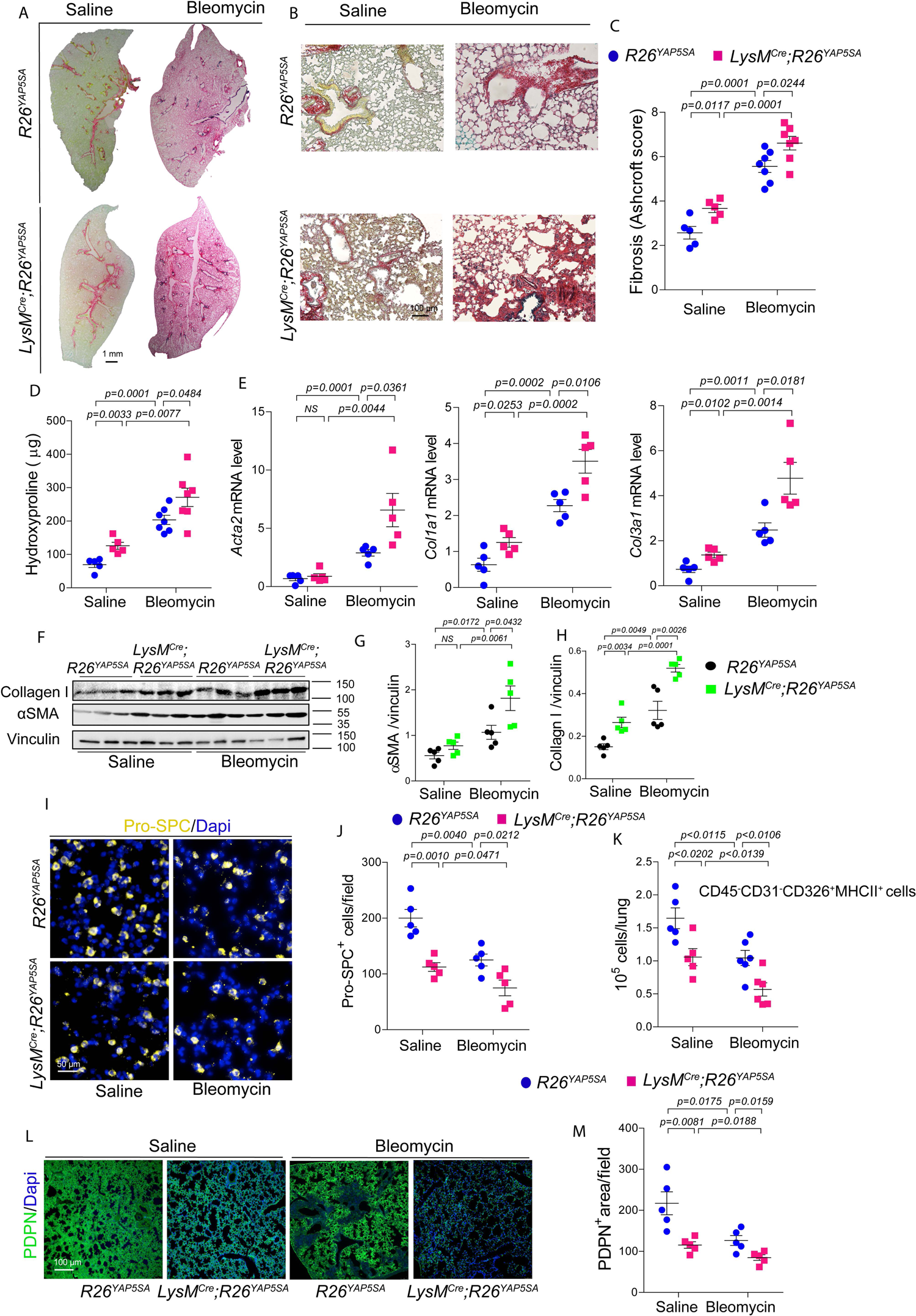
Macrophage-specific *Yap* overexpression augments bleomycin-induced lung fibrosis in mice. (A-C) Sirius Red/Fast Green staining and quantification (presented as Ashcroft score) on lung sections from *R26^YAP5SA^*and *LysM^Cre^;R26^YAP5SA^* mice (n = 5-7) treated with saline or bleomycin for 14 days. (D) Quantification of hydroxyproline content using lung tissue from *R26^YAP5SA^* and *LysM^Cre^;R26^YAP5SA^* mice (n = 5-7) treated with saline or bleomycin for 14 days. (E) Real-time qPCR for profibrotic genes *Acta2, Col1a1,* and *Col3a1* on lung tissue from *R26^YAP5SA^* and *LysM^Cre^;R26^YAP5SA^* mice (n = 5) treated with saline or bleomycin for 14 days. (F-H) Immunoblot analysis and quantification of collagen I and αSMA protein using lung tissue from *R26^YAP5SA^*and *LysM^Cre^;R26^YAP5SA^* mice (n = 5) treated with saline or bleomycin for 14 days. (I-J) Immunostaining and quantification of Pro-SPC^+^ AT2-cells in *R26^YAP5SA^ versus LysM^Cre^;R26^YAP5SA^* mice lung (n = 5) treated with saline or bleomycin for 14 days. (K) Flow cytometry analysis to determine the changes in the number of AT2-cells identified as CD45^−^CD31^−^CD326^+^MHCII^+^ in *LysM^Cre^;R26^YAP5SA^* lungs compared to *R26^YAP5SA^* lungs (n = 5) after saline or bleomycin treatment for 14 days. (L-M) Immunostaining and quantification of PDPN^+^ AT1-cells in *LysM^Cre^;R26^YAP5SA^* lungs compared to *R26^YAP5SA^* lungs (n = 5) after saline or bleomycin treatment for 14 days. The data are represented as the mean ± SEM; comparison by two-tailed unpaired t-test. *, *p* < 0.05; **, *p* < 0.01; ***, *p* < 0.001; NS, not significant.

### YAP overexpression augments bleomycin-induced alveolar epithelial cell damage

To understand whether YAP overexpression modulates the alveolar epithelial regeneration *in vivo*, *R26^YAP5SA^* and *LysM^Cre^;R26^YAP5SA^*mice were exposed to either saline or bleomycin for 14 days. Lung tissues were collected to perform the immunohistochemistry, flow cytometry and qRT-PCR analysis. In saline treated group, qRT-PCR analysis showed a significant downregulation of AT2 marker *Sftpc* expression in *LysM^Cre^;R26^YAP5SA^*lung compared to control. Bleomycin administration further dampened the *Sftpc* expression in both group; while in *LysM^Cre^;R26^YAP5SA^*lung, *Sftpc* level reached to baseline (Supplementary figure 7A). Next, using immunofluorescence we determined the AT2 cell numbers in *R26^YAP5SA^* and *LysM^Cre^;R26^YAP5SA^*. Compared to saline treated group, bleomycin treated lung showed a profound damage of epithelial cells in the lung as indicated by reduced the number of pro-SPC-expressing AT2 cells and YAP overexpression caused a substantial loss of AT2 cells in both saline and bleomycin-treated group (Figure 6I-J). Flow cytometry analysis was correlated with these observations, as we have seen a significant decrease in the number of AT2 cells identified by CD45^−^CD31^−^CD326^+^MHCII^+^ population in *LysM^Cre^;R26^YAP5SA^* lung compared with *R26^YAP5SA^* lung (Figure 6K). Similar to AT2, AT1 cells were also negatively affected after bleomycin-induced injury. In saline treated groups, no alteration were detected between *R26^YAP5SA^* and *LysM^Cre^;R26^YAP5SA^* lung, whereas an intense loss of AT1 cells was observed in *LysM^Cre^;R26^YAP5SA^* lung compared to *R26^YAP5SA^* lung post-injury, indicated by the PDPN positive areas (Figure 6L-M).. Consistently, expression of another AT1 marker *Cav1* was downregulated after bleomycin and it was further reduced in *LysM^Cre^;R26^YAP5SA^* lung compared to *R26^YAP5SA^* lung. However, saline treatment did not affect the expression of *Cav1* between these groups (Supplementary figure 7B). These findings suggests a critical role macrophage-YAP-mediated epithelial regeneration after injury.

### Blocking of CCL2 using CCL2 neutralizing antibody suppresses YAP-mediated pulmonary fibrosis after bleomycin injury

Macrophage infiltration plays a crucial role in the pathogenesis of IPF/PF. After lung injury, macrophage trafficking to inflammatory sites is mostly regulated by the inflammatory chemokine CCL2, also known as monocyte chemoattractant protein-1 (MCP-1), pathway (Moore, Paine et al., 2001, Okuma, Terasaki et al., 2004). Elevated expression of CCL2 have been described in preclinical models (Moore et al., 2001, Smith, Strieter et al., 1994) and in patients with IPF as well as in inflammatory lung diseases (Antoniades, Neville-Golden et al., 1992, Iyonaga, Takeya et al., 1994, Rose, Sung et al., 2003, Sefik, Qu et al., 2022, Suga, Iyonaga et al., 1999). Blocking of CCL2 with neutralizing antibody or genetic deletion of its receptor CCR2 attenuated PF (Gharaee-Kermani, McCullumsmith et al., 2003, Moore et al., 2001, Okuma et al., 2004), suggesting a potential of CCL2 in lung fibrogenesis. In the current study, we sought to examine whether the expression and function of CCL2 was alerted due to the macrophage-specific deletion of *Yap/Taz* or overexpression of *Yap*. Using qRT-PCR analysis we have shown that deficiency of *Yap/Taz* reduced the mRNA expression of *Ccl2* in bleomycin-injured lung (Figure 2D); likewise expression of *Ccl2* was also attenuated in Mo-AMs (Figure 3E) but not in IMs (Figure 3F). Overexpression of *Yap* augmented *Ccl2* gene expression in lung (Figure 5D) as well as in sorted Mo-AMs after bleomycin injury (Figure 5H). Immunoblot analysis revealed reduced protein expression of CCL2 in *dKO* lung compared to respective control after bleomycin (Figure 7A); however CCL2 expression was increased in *LysM^Cre^;R26^YAP5SA^* lung compared to *R26^YAP5SA^* lung both in saline and bleomycin treated animals at 14 days (Figure 7B). These observation suggesting YAP/TAZ may regulate lung fibrosis through the activation of CCL2. We next found that ∼5 kb of genomic sequence upstream of the first exon of *Ccl2* promoter contains multiple predicted TEAD binding motifs. To understand whether YAP and TAZ binds to the promoter of *Ccl2*, we performed chromatin immunoprecipitation (ChIP) assay using bone-marrow derived macrophages (BMDMs) treated with pro-inflammatory stimuli by lipopolysaccharide (LPS). We found that YAP and TAZ-immunoprecipitations were enriched with *Ccl2* promoter DNA (Figure 7C). To demonstrate the YAP-mediated role CCL2 in PF, we have utilised *LysM^Cre^;R26^YAP5SA^* animals and treated the mice with IgG or CCL2 neutralizing antibody after bleomycin administration for 14 days (Figure 7D). Blocking of CCL2 with neutralizing antibody reduced CD68+ macrophage infiltration in *LysM^Cre^;R26^YAP5SA^*fibrotic lung (Figure 7E). Flow cytometry analysis revealed a decrease in Mo-AMs population while the TR-AMs and IMs population were unaffected due to CCL2 neutralization (Figure 7F). Sirius red staining showed a lower fibrotic Ashcroft score in *LysM^Cre^;R26^YAP5SA^* + anti-CCL2 lung compared to IgG treated lung (Figure 7G-H). Likewise, collagen III protein expression was attenuated in response to anti-CCL2 treatment as identified by immunoblot (Figure 7I). Gene expression analysis revealed a downregulation of proinflammatory and profibrotic genes such as *Il6*, *Il1β*, *Acta2*, *Col1a1*, and *Ctgf* in anti-CCL2 treated *LysM^Cre^;R26^YAP5SA^*lung (Figure 7J). In addition, we observed an improved in the number of pro-SPC+ (as determined by immunofluorescence analysis) and CD45^−^CD31^−^CD326^+^MHCII^+^ (as determined by the flow cytometry analysis) AT2-cells in *LysM^Cre^;R26^YAP5SA^* + anti-CCL2 lung (Figure 7K-L). As similar to AT2 cells, the AT1 cells, as defined the PDPN+ area, were also protected due to anti-CCL2 administration in bleomycin treated *LysM^Cre^;R26^YAP5SA^*lung (Figure 7M). These data indicate that YAP activation in macrophages causes augmented inflammation, fibrosis and epithelial damage possibly due to CCL2 mediated proinflammatory macrophage infiltration.

**Figure 7:**
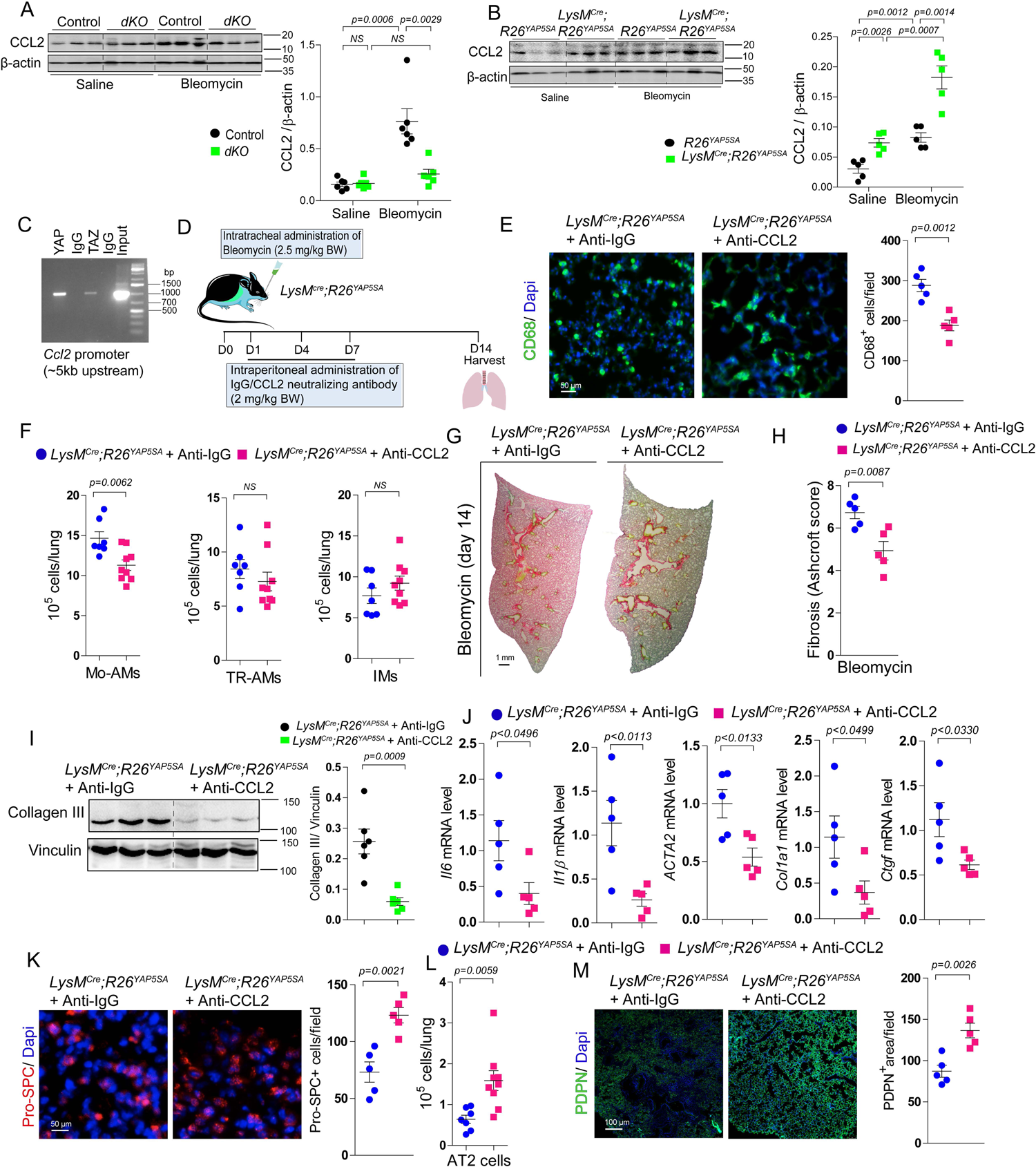
YAP/TAZ modulates CCL2 expression and neutralizing of CCL2 attenuates YAP-induced pulmonary fibrosis after bleomycin-injury. (A) Immunoblot analysis and quantification of CCL2 protein using lung tissue from control and *dKO* mice (n = 5-6) treated with saline or bleomycin for 14 days. (B) Immunoblot analysis and quantification of CCL2 protein using lung tissue from *R26^YAP5SA^* and *LysM^Cre^;R26^YAP5SA^* mice (n = 5) treated with saline or bleomycin for 14 days. (C) ChIP assay performed on *Ccl2* promoter using chromatin from wildtype BMDMs exposed to LPS (100 ng/mL) using IgG and YAP or TAZ antibody. (D) Graphical presentation of experimental design; *LysM^Cre^;R26^YAP5SA^*mice were challenged to intratracheal administration of bleomycin (as 2.5 mg/kg body weight) with following intraperitoneal injection of CCL2 neutralizing antibody or IgG (as 2mg/kg body weight) as indicated time points. (E) Immunostaining and quantification of CD68 positive macrophage infiltration in *LysM^Cre^;R26^YAP5SA^* mice lung (n = 5) treated with bleomycin followed IgG or CCL2 neutralizing antibody treatment for 14 days. (F) Quantification of flow cytometry analysis to sort Mo-AMs (identified as CD45^+^Ly6G^-^CD64^+^CD11b^low^Siglec-F^low^), TR-AMs (identified as CD45^+^Ly6G^-^ CD64^+^CD11b^low^Siglec-F^high^) and IMs (CD45^+^Ly6G^-^CD64^+^CD11b^high^Siglec-F^-^) using lung from *LysM^Cre^;R26^YAP5SA^*mice (n = 7-9) treated with bleomycin followed IgG or CCL2 neutralizing antibody treatment for 14 days. (G-H) Sirius Red/Fast Green staining and quantification (presented as Ashcroft score) on lung sections from *LysM^Cre^;R26^YAP5SA^*mice (n = 5) treated with bleomycin following IgG or CCL2 neutralizing antibody treatment for 14 days. (I) Immunoblot analysis and quantification of collagen III protein in lung tissue from *LysM^Cre^;R26^YAP5SA^* mice (n = 5) treated with bleomycin followed IgG or CCL2 neutralizing antibody treatment for 14 days. (J) mRNA analysis of *Il6*, *Il1β*, *Acta2, Col1a1* and *Ctgf* on lung tissue from *LysM^Cre^;R26^YAP5SA^* mice (n = 5) treated with bleomycin followed IgG or CCL2 neutralizing antibody treatment for 14 days. (K) Immunostaining and quantification of Pro-SPC^+^ AT2-cells in *LysM^Cre^;R26^YAP5SA^* mice (n = 5) treated with bleomycin followed IgG or CCL2 neutralizing antibody treatment for 14 days. (L) Flow cytometry analysis to determine the changes in the number of AT2-cells identified as CD45^−^CD31^−^CD326^+^ MHCII^+^ in *LysM^Cre^;R26^YAP5SA^* lungs (n = 7-9) treated with bleomycin followed IgG or CCL2 neutralizing antibody treatment for 14 days. (M) Immunostaining and quantification of PDPN^+^ AT1-cells in *LysM^Cre^;R26^YAP5SA^* lungs (n = 5) treated with bleomycin following IgG or CCL2 neutralizing antibody treatment for 14 days. The data are represented as the mean ± SEM; comparison by two-tailed unpaired t-test. *, *p* < 0.05; **, *p* < 0.01; ***, *p* < 0.001; NS, not significant.

### MBD2 is activated in lung macrophages in bleomycin-treated mice and act as downstream target of YAP during profibrotic macrophage polarization

Methyl-CpG–binding domain 2 (MBD2), is a DNA methylation reader, has been recently shown to promote the resting macrophages to polarize into profibrotic macrophages during bleomycin-induced PF (Wang, Zhang et al., 2021b). MBD2 was also found to be largely expressed in macrophages of patient’s lungs with IPF as well as in mice lungs derived from bleomycin-induced PF (Wang et al., 2021b). Therefore, we sought to explore whether YAP/TAZ are involved to regulate the expression and function of MBD2 in macrophages during PF. To confirm the expression of MBD2 in macrophages we utilised the lung sections from PF patients and mice lung from bleomycin-induced PF with their respective controls. As expected, the presence of MBD2 in macrophages as detected by CD68 and MBD2-double-positive cells (CD68^+^ MBD2^+^ cells) were highly detected in the PF patients lung compared to control adult human lung (Figure 8A). Next we analyzed the presence of CD68^+^ MBD2^+^ cells in mice PF model. Bleomycin instillation increased the number of CD68^+^ MBD2^+^ cells compared to saline treatment (Figure 8B). Co-immunostaining of MBD2 with YAP and TAZ in CD68^+^ macrophages suggested that MBD2 was predominantly colocalized with YAP/TAZ in macrophages during PF in human and mice (Supplementary figure 8A-B). Next we observed whether macrophage-specific deletion of *Yap/Taz* or overexpression of *Yap* affect the MBD2 expression in bleomycin-induced mice lung. Interestingly, genetic loss of *Yap/Taz* reduced the MBD2 protein expression, while overexpression of *Yap* augmented its protein level after bleomycin injury compared to saline treated uninjured lung (Figure 8C-F). Immunostaining confirmed the immunoblot data, deletion of *Yap/Taz* significantly attenuated CD68^+^ MBD2^+^ macrophage numbers; however *Yap* overexpression increased the number of CD68^+^ MBD2^+^ cells in lung challenged to bleomycin (Supplementary figure 8C-D). Consequently, the protein expression of profibrotic macrophage marker ARG1 was also attenuated in *dKO* lung, whereas *LysM^Cre^;R26^YAP5SA^*lung promoted ARG1 expression (Figure 8C-F), suggesting a possible correlation between the YAP/TAZ and MBD2 in macrophage polarization during bleomycin-induced PF.

**Figure 8:**
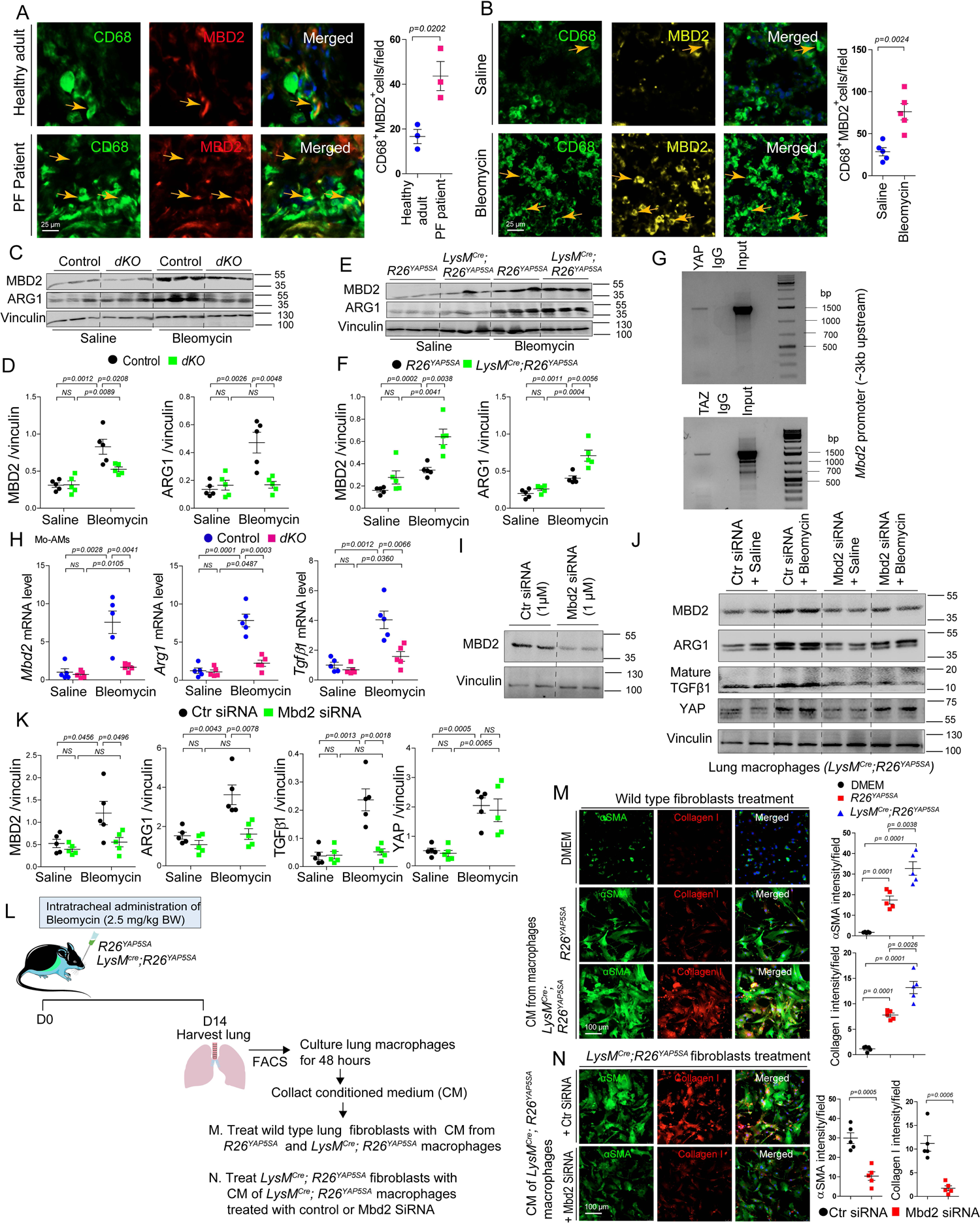
Loss of *Yap/Taz* attenuates MBD2 expression in mice fibrotic lung and MBD2 act as downstream of YAP during profibrotic macrophage polarization. (A) MBD2 is highly expressed in CD68+ lung macrophages of pulmonary fibrotic patient compared to normal adult human (healthy adult) subject (n = 3). (B) MBD2 is highly expressed in CD68+ lung macrophages after bleomycin treatment compared to saline treatment in wildtype mice (n = 5). (C-D) Immunoblot analysis and quantification of MBD2 and ARG1 protein using lung tissue from control and *dKO* mice (n = 5) treated with saline or bleomycin for 14 days. (E-F) Immunoblot analysis and quantification of MBD2 and ARG1 protein using lung tissue from *R26^YAP5SA^* and *LysM^Cre^;R26^YAP5SA^*mice (n = 5) treated with saline or bleomycin for 14 days. (G) ChIP assay performed on *Mbd2* promoter using chromatin from wildtype BMDMs exposed to LPS (100 ng/mL) using IgG and YAP or TAZ antibody. (H) Real-time qPCR for *Mbd2*, *Arg1* and *Tgfβ1* on sorted Mo-AMs from control and *dKO* mice (n = 5) treated with saline or bleomycin for 14 days. (I) Immunoblot analysis to confirm the knockdown efficiency of MBD2 from knockdown experiment with *Mbd2* siRNA or control siRNA using primary cultured total lung macrophages (CD45^+^Ly6G^-^ CD64^+^CD11b^+^). (J-K) Immunoblot analysis and quantification of MBD2, ARG1, mature-TGFβ1 and YAP using total lung macrophages (CD45^+^Ly6G^-^CD64^+^CD11b^+^) isolated by flow cytometry and subsequently cultured derived from *LysM^Cre^;R26^YAP5SA^*mice lung (n = 5) treated with bleomycin or saline to perform the knockdown experiment with *Mbd2* siRNA or control siRNA. (L) Graphical presentation of experimental design; *R26^YAP5SA^* and *LysM^Cre^;R26^YAP5SA^*mice were challenged to intratracheal administration of bleomycin (as 2.5 mg/kg body weight) and harvested the total lung macrophages and cultured to collect the conditioned medium (CM). (M) Immunofluorescence staining and quantification of αSMA and collagen I using wildtype lung (n = 5) fibroblasts treated with either DMEM or CM collected from *R26^YAP5SA^* and *LysM^Cre^;R26^YAP5SA^*lung macrophages challenged to bleomycin-injury. (N) Immunofluorescence staining and quantification of αSMA and collagen I using *LysM^Cre^;R26^YAP5SA^*lung (n = 5) fibroblasts treated with CM collected from *LysM^Cre^;R26^YAP5SA^* lung macrophages challenged to bleomycin-injury followed by *Mbd2* siRNA or control siRNA exposure. The data are represented as the mean ± SEM; comparison by two-tailed unpaired t-test. *, *p* < 0.05; **, *p* < 0.01; ***, *p* < 0.001; NS, not significant.

Next we analyzed the ∼3 kb of genomic sequence upstream of the first exon of MBD2 which revealed multiple predicted TEAD-binding sequences to its promoter. To determine whether YAP and TAZ binds to the promoter of *Mbd2*, we performed ChIP experiment using BMDMs treated with LPS and found that YAP and TAZ-immunoprecipitated products were enriched with *Mbd2* promoter DNA (Figure 8G), suggesting YAP/TAZ may regulate MBD2-mediated macrophage function. Indeed, qRT-PCR analysis on sorted Mo-AMs revealed that *Mbd2* expression was downregulated together with profibrotic macrophage gene *Arg1* and *Tgfβ1* in *dKO* lung after bleomycin administration (Figure 8H). To dissect the YAP-mediated role of MBD2 in macrophage further we have used the *LysM^Cre^;R26^YAP5SA^* mice. We treated the *LysM^Cre^;R26^YAP5SA^*mice with bleomycin or saline and harvested the total lung macrophages (CD45^+^Ly6G^-^CD64^+^CD11b^+^) using flow cytometry and subsequently cultured the cells to perform the knockdown experiment with *Mbd2* siRNA or control siRNA (Figure 8I-K). We also harvested the conditioned medium (CM) from the cultured lung macrophages treated with *Mbd2* siRNA or control siRNA. The knockdown efficiency of *Mbd2* in primary lung macrophages was confirmed by immunoblot analysis (Figure 8I). Compared to saline, bleomycin augmented the protein levels of ARG1, mature-TGFβ1, and YAP in primary lung macrophages. Interestingly, knockdown of *Mbd2* significantly abrogated the bleomycin-induced ARG1 and mature TGFβ1 protein expression in lung macrophages, whereas the levels of YAP protein was not altered (Figure 8J-K). These observation suggesting MBD2 acts as downstream of YAP to induce profibrotic macrophage polarization during bleomycin-induced PF.

It is well established that the secretome of macrophage containing the profibrotic mediators is crucial to activate the neighbouring cells including fibroblasts in PF (Lis-Lopez, Bauset et al., 2021). To demonstrate whether YAP mediated function of MBD2 in macrophage affects the profibrotic events in fibroblasts, we collected the CM from the cultured lung macrophages harvested from *R26^YAP5SA^* and *LysM^Cre^;R26^YAP5SA^* lung treated with bleomycin (Figure 8L). We then treated the wildtype lung fibroblasts with normal DMEM as control or CM of *R26^YAP5SA^* and *LysM^Cre^;R26^YAP5SA^*lung macrophage and observed the myofibroblasts formation and collagen I production. In comparison to DMEM control CM of *R26^YAP5SA^* macrophages significantly induces αSMA and collagen I expression and their expression was further augmented when we treated the fibroblasts with CM of *LysM^Cre^;R26^YAP5SA^*macrophages (Figure 8M), suggesting secretome from YAP overexpressing macrophage is responsible to stimulate the fibroblasts activation into myofibroblasts formation. To determine whether knockdown of *Mbd2* in *LysM^Cre^;R26^YAP5SA^*lung macrophages affect YAP-mediated profibrotic events in fibroblasts, we treated the *LysM^Cre^;R26^YAP5SA^* lung fibroblasts with CM of *LysM^Cre^;R26^YAP5SA^* lung macrophages treated with *Mbd2* siRNA or control siRNA. Indeed, *Mbd2* siRNA containing CM was able to inhibit the induction αSMA and collagen I production by *LysM^Cre^;R26^YAP5SA^* fibroblasts (Figure 8N), indicating YAP stimulates the profibroitc function through MBD2 in bleomycin-induced PF.

## Discussion

IPF/PF is resulting from the consequence of number of deleterious insult including the infiltration of inflammatory monocytes/macrophages into the injured lung. The numbers of lung macrophages are prominent both in patients with IPF (Byrne, Maher et al., 2016, Schwartz, Helmers et al., 1991), and in preclinical models of PF (Osterholzer, Olszewski et al., 2013, Redente, Keith et al., 2014). The existence of multiple subset of lung macrophages in PF/IPF is well-established, however their relative contribution and regulation in fibrogenesis is incompletely understood. In addition, the mechanism of macrophage-mediated proinflammatory response that impact fibrogenesis and its potential as therapeutic target in IPF/PF is poorly characterized. In the current study, we demonstrate that both Mo-AMs and IMs population contributes to the development of bleomycin-induced fibrosis, although the fibrogenic role of Mo-AMs was more robust compared to IMs. We found that YAP and TAZ are activated in lung macrophages of patients diagnosed with PF and in mice followed bleomycin-induced injury. Macrophage-specific conditional deletion of *Yap/Taz* decreased the number of Mo-AMs and IMs population in bleomycin-treated lung. As a consequence, genetic loss of *Yap/Taz* impaired the development of fibrosis by deteriorating macrophage-mediated inflammatory response and improved repair capacity of alveolar epithelial cells AT1 and AT2 after bleomycin-induced injury. In contrast, overexpression of *Yap* in macrophages augmented the Mo-AMs population, promoted the inflammatory response to the development of fibrosis and worsened the alveolar epithelial damage during post-bleomycin injury.

Macrophages in lung are comprised of heterogeneous cell population known to be involved in tissue homeostasis, inflammation, and numerous pathologies including fibrogenesis. Several preclinical studies demonstrated the contribution of macrophages in inflammation-induced fibrosis and distinct subpopulations of macrophages differentially involved to PF (Chakarov et al., 2019, Cui et al., 2020, Gibbons, MacKinnon et al., 2011, Joshi et al., 2020, McCubbrey et al., 2018, Misharin et al., 2017, Osterholzer et al., 2013, Singh, Chakraborty et al., 2022). However, the regulatory mechanisms and proinflammatory/profibrotc behaviour of distinct subpopulations in PF remain poorly understood. Here, we demonstrate that YAP/TAZ are critical regulator of macrophage recruitments and macrophage-mediated inflammatory response in mice lung after bleomycin-induced injury. Studies evident that aberrant activation of profibrotic Mo-AMs and their interaction with fibroblasts is a major determinant for the development of lung fibrogenesis (Aran, Looney et al., 2019, Gharaee-Kermani et al., 2003, Joshi et al., 2020, Misharin et al., 2017, Redente et al., 2014, Singh et al., 2022). After lung injury, AMs population are increasing and studies indicate that circulating monocytes are differentiating into macrophages, defined as Mo-AMs, which are playing the profibrotic role to induce PF. Studies suggest that depletion of circulating monocytes using *Ccr2−/−* mice or using liposomal clodronate significantly decreased the severity of PF (Moore et al., 2001). Likewise, targeted depletion of Mo-AMs through necroptosis during their differentiation by using *Casp8^flox/flox^* mice ameliorated the lung fibrosis (Joshi et al., 2020, Misharin et al., 2017); however depletion of TR-AMs did not contribute to induce fibrosis (Misharin et al., 2017). Studies also illustrate that depletion of Mo-AMs reduces the production of profibrotic component (Joshi et al., 2020, McCubbrey et al., 2018), implicating the crucial role of Mo-AMs in the development of fibrosis. Consistent to these findings, our study also demonstrate an increased profibrotic Mo-AMs (CD45^+^Ly6G^-^CD64^+^CD11b^low^Siglec-F^low^) population during bleomycin-induced lung fibrosis and this augment was prominent after 14 days of bleomycin instillation. Myeloid-specific genetic deletion of *Yap/Taz* depleted the number of Mo-AMs and as a consequence the gene expression of proinflammatory/profibrotic cytokines and chemokines such as *Il6*, *Il1β*, *Ccl2,* and *Cx3cr1* were reduced in the bleomycin-injured lung. By overexpressing *Yap* in macrophages we were able to demonstrate an elevated number of Mo-AMs with the upregulated gene expression profiling of *Il6*, *Il1β*, *Ccl2,* and *Cx3cr1* by this macrophages after bleomycin treatment. Not like AMs, the number of IMs population (identified as CD45^+^Ly6G^-^ CD64^+^CD11b^high^Siglec-F^-^) were substantially decreased at day 7, whereas they moderately increased at day 14 post-bleomycin administration and knockout of *Yap/Taz* reduced the numbers of IMs at day 14 bleomycin injury. Likewise, loss of *Yap/Taz* reduced the expression of *Il6*, *Il1β*, and *Cx3cr1*, but not *Ccl2* expression. We also observed another critical difference between the Mo-AMs and IMs. As such, there was a high-fold upregulation of *Il6*, *Il1β*, *Ccl2,* and *Cx3cr1* genes in Mo-AMs at day 14 post-bleomycin injury; however these genes were moderately increased in IMs, suggesting the contribution Mo-AMs in the development of PF is more potent compared to IMs although both population involved to express profibrotic components after injury. Our findings revealed the gene expression data from mix population of IMs, which presenting one of the limitation of our study. Since, a recent study reported two distinct IMs lineage from monocyte derived tissue resident macrophages identified as Lyve1^low^MHCII^high^ or Lyve1^high^MHCII^low^. Depletion of Lyve1^high^MHCII^low^ IMs exacerbated the bleomycin-induced PF (Chakarov et al., 2019), indicating functional diversity of distinct subset of IMs is crucial in the pathogenesis of PF. Further study will be needed to distinguish the differential role of YAP/TAZ toward the regulation of subpopulation of IMs. Moreover, depletion of TR-AMs did not contribute to induce fibrosis (Misharin et al., 2017); in the line, our study showed the number of TR-AMs was unaffected due to deletion of *Yap/Taz* or over expression of *Yap* in bleomycin-injured lung. Together, our results suggest that YAP/TAZ are important regulator of Mo-AMs infiltration and secretion of fibrogenic factors by these cells.

Inflammasome activation and its mediated inflammatory response are crucial during lung inflammation, since blocking of the NLRP3 infammasome pathway has been reported to be beneficial for the prevention of chronic lung pathophysiology including IPF (Jager, Seeliger et al., 2021, Lasithiotaki, Giannarakis et al., 2016, Sefik et al., 2022). Recent studies revealed that YAP promotes the activation of NLRP3 inflammasome and deficiency of YAP in macrophages significantly reduced the LPS-induced systemic inflammation and (Wang, Zhang et al., 2021a). Currently, it is not known how deficiency of *Yap/Taz* in lung macrophages affects their function and thereby regulates infammasome activity and inflammation in lung after injury. Here we demonstrate that *Yap/Taz* deficiency reduces the production of caspase-1 and IL-1β by lung macrophages after bleomycin-induced injury, suggesting that loss of *Yap/Taz* abrogates the proinflammatory events through NLRP3 inflammasome activation process in macrophages, which is important for caspase-1-mediated maturation of IL-1β during inflammasome activation to induce inflammatory response (Kayagaki et al., 2011).

Mo-AMs are crucial drivers of lung fibrosis (Chakarov et al., 2019, Misharin et al., 2017). Depletion of lung macrophages or circulating monocytes prevented the PF, however lung macrophages also exhibiting the resolution-promoting role during in PF (Cui et al., 2020, Gibbons et al., 2011, Misharin et al., 2017). The contribution of monocytes and macrophages is controversial, but phase-dependent during wound healing or fibrosis. Therefore targeting the regulatory mechanism of macrophage phenotype could be essential to prevent the PF. Previously, we demonstrated that macrophage specific deficiency of *Yaz/Taz* resulted in improved cardiac remodeling and function after myocardial infarction due to reduced cardiac fibrosis (Mia et al., 2020). Consistently, in the current study we also found that *Yap/Taz* inactivation in macrophages impaired fibrosis as described by the reduced αSMA+ myofibroblast formation and collagen synthesis in bleomycin-induced lung. Similarly, overexpression of YAP augmented the fibrotic response in the lung followed bleomycin. We also investigated the regenerative capacity of the damaged lung after bleomycin-injury, since impairment of alveolar AT2 cells has been described to guides the progression of IPF/PF (Selman & Pardo, 2020, Sisson et al., 2010). In lung alveoli, AT2 cells function as resident stem cells which can proliferate and differentiate into AT1 cells required to regenerate the development of new alveoli due to pulmonary damage after injury (Adamson & Bowden, 1974, Barkauskas et al., 2013). A recent study demonstrate that deficiency of *Yap/Taz* in AT2 cells augmented inflammatory responses in the lung which resulted a delay in alveolar epithelial regeneration during bacterial pneumonia (LaCanna, Liccardo et al., 2019). Our findings on macrophage-specific *Yap/Taz* double-knockout or *Yap*-overexpression provided the new insights in alveolar regeneration after bleomycin-injury. We showed that lacking of *Yap/Taz* in macrophage increased the number of AT2 cells (identified as CD45^−^CD31^−^CD326^+^MHCII^+^) as well as PDPN-positive AT1 cells at 14 day post-injury. In contrast, Yap overexpression depletes the both AT2 and AT1 cells at post-bleomycin injury; suggesting cell-specific function of YAP/TAZ is crucial to regulate the AT1 and AT2-cells mediated alveolar epithelial repair after injury.

It is increasingly reported that infiltrating macrophages with proinflammatory/profibrotic nature into the injured lung trigger the development of fibrosis (Joshi et al., 2020, Misharin et al., 2017, Moore et al., 2001, Redente et al., 2014, Singh et al., 2022). Therefore, targeting the macrophage recruitment pathway could be a beneficial approach for the treatment of PF. The inflammatory chemokines CCL2-CCR2 pathway play a key role to drive the macrophage immigration after lung injury (Moore et al., 2001, Okuma et al., 2004). CCR2 is required to promote the lung fibrosis; however deficiency of CCR2 systemically in mice resulted in a defective macrophage migration (Moore et al., 2001), reduced profibrotic cytokine expression and impaired myofibroblasts formation (Gharaee-Kermani et al., 2003). Higher expression of CCL2 was also associated with fibrosis both in preclinical models (Moore et al., 2001, Smith et al., 1994) and in patients with inflammatory lung diseases including IPF (Antoniades et al., 1992, Iyonaga et al., 1994, Rose et al., 2003, Sefik et al., 2022, Suga et al., 1999). Inhibition of CCL2 with neutralizing antibody or genetic ablation of its receptor CCR2 attenuated macrophage recruitment and improved lung fibrosis in mice (Gharaee-Kermani et al., 2003, Moore et al., 2001, Okuma et al., 2004), indicating CCL2-CCR2 signaling is involved in the pathogenesis of PF/IPF by regulating fibrogenic cytokine expression and fibroblast activation. Using ChIP analysis, we demonstrate that YAP/TAZ bind to the *Ccl2* promoter in macrophages, which suggesting YAP/TAZ may regulate CCL2 functions. Indeed, we found that deletion of *Yap/Taz* attenuated the *Ccl2* and its receptor *Ccr2* expression in bleomycin injured lung. *Ccl2* gene expression was also downregulated in Mo-AMs not in IMs, although we were unable to detect *Ccr2* expression in these cells. Consistently, the expression *Ccl2* and *Ccr2* were promoted in macrophage-specific *Yap*-overexpressing lung; as expected *Ccl2* gene level was elevated in isolated Mo-AMs, but not in IMs. Blocking of with CCL2 neutralizing antibody injected in *LysM^Cre^;R26^YAP5SA^*mice, we observed an improved fibrosis with alveolar epithelial repair as consequence of reduced CD68^+^ macrophage infiltration, lesser number of infiltrating Mo-AMs, and impaired proinflammatory response after bleomycin-induced lung injury. These data suggesting YAP/TAZ may regulate the CCL2-mediated macrophage recruitment and profibrotic response induced by fibrogenic cytokines released from Mo-AMs.

Profibrotic macrophage-polarization mediators could also be a fruitful targets to prevent the progression of fibrosis. Recent studies implicated the involvements of MBD2 in organ fibrosis including PF (Ai, Pan et al., 2022, Wang, Zhang et al., 2022, Wang et al., 2021b), mostly described its role as a DNA methylation reader and partner of Nucleosome Remodelling and Deacetylation (NuRD) complex that regulating cell fate transitions and differentiation (Wood & Zhou, 2016), which induces the polarization of resting macrophages into profibrotic macrophages in bleomycin-induced mice (Wang et al., 2021b). Therefore, we sought to explore whether YAP/TAZ are involved to regulate the expression and function of MBD2 in macrophages during PF. MBD2 has been shown to be largely expressed in macrophages of patients lung with IPF and in mice lung followed bleomycin-induced PF (Wang et al., 2021b). Deficiency of *Mbd2* in macrophages reported to be protective against bleomycin induced PF in mice. *Mbd2* deficiency in macrophages notably reduced TGF-β1 production and attenuated ARG1^+^ profibrotic macrophage accumulation in the lung after bleomycin-induced injury (Wang et al., 2021b). Like macrophages, deficiency of *Mbd2* in fibroblasts or myofibroblasts also protected the mice from bleomycin-induced PF by attenuating the activation of fibroblast through interrupting TGF-β1-dependent Smad2/3 pathway (Wang et al., 2022). Consistent to these studies, our finding incorporating the essential role of YAP/TAZ in regulating MBD2 functions towards macrophage polarization and fibroblasts activation in PF. Macrophage-specific *Yap/Taz* deficiency reduced the protein expression of MBD2, while Yap overexpression promoted its protein level in mice lung followed bleomycin induction. Using ChIP assay on BMDMs, we assessed that YAP/TAZ bind to the *Mbd2* promoter in macrophages. Depletion of *Yap/Taz* in macrophages downregulated *Mbd2* expression on sorted Mo-AMs together with profibrotic macrophage genes *Arg1* and *Tgfβ1*, suggesting a potential correlation of YAP/TAZ with MBD2 to induce profibrotic macrophage polarization. Indeed, knockdown experiment of *Mbd2* on sorted total macrophages (CD45^+^Ly6G^-^CD64^+^CD11b^+^) from *LysM^Cre^;R26^YAP5SA^* lung using primary cell-specific siRNA abrogated the bleomycin-induced protein levels of ARG1, mature-TGFβ1, while YAP expression was unaffected due to *Mbd2* knockdown; indicating MBD2 acts as downstream of YAP to simulate profibrotic macrophage polarization and profibrotic cytokine TGFβ1 secretion in bleomycin-induced lung. Earlier studies suggested compelling evidence that release of TGFβ1, the main driver of fibrotic diseases, by profibrotic M2 macrophages contributes to the transition of fibroblast into collagen-producing myofibroblasts formation (Boutanquoi, Burgy et al., 2020, Hou, Shi et al., 2018). Therefore, we addressed whether YAP-MBD2 mediated macrophage-fibroblasts crosstalk in our study affects myofibroblasts induction and collagen production. Indeed, we found that treatment of bleomycin-induced *LysM^Cre^;R26^YAP5SA^* lung fibroblasts with CM of *LysM^Cre^;R26^YAP5SA^* lung macrophages augmented αSMA+ myofibroblast formation and collagen I expression, however CM of *LysM^Cre^;R26^YAP5SA^*lung macrophages treated with *Mbd2* siRNA significantly inhibited these profibrotic events in lung fibroblasts. These observations proposed a defective TGFβ1 secretion due to *Mbd2* knockdown in *LysM^Cre^;R26^YAP5SA^*lung macrophages displaying the lower expression of αSMA and collagen I. Together, these findings supported the evidence that YAP regulates the macrophage-fibroblasts crosstalk through MBD2 that induces the transdifferentiation of fibroblasts into myofibroblasts in bleomycin-induced PF.

In summary, *Yap/Taz* are potent regulator of macrophage infiltration after bleomycin-induced injury (Figure 9). Macrophage-specific genetic depletion of *Yap/Taz* impaired the immigration of Mo-AMs which resulting a defective macrophage-mediated inflammatory response and improved pulmonary fibrosis in bleomycin-treated lung. However, overexpression of *Yap* in macrophages augmented the Mo-AMs recruitment, stimulated the proinflammatory response and worsened the fibrosis progression at post-bleomycin injury. Mechanistically, we found that YAP/TAZ regulate bleomycin-induced lung fibrosis through the activation of CCL2 that drives macrophage recruitment and neutralizing of CCL2 with antibody abrogated YAP-mediated profibrotic outcomes. We also identified that YAP regulates the macrophage-fibroblasts crosstalk through MBD2-TGFβ1-axis to induce the differentiation of fibroblasts into myofibroblasts and MBD2 acts as downstream of YAP to induce ARG1+ profibrotic macrophage polarization during bleomycin-induced PF.

**Figure 9:**
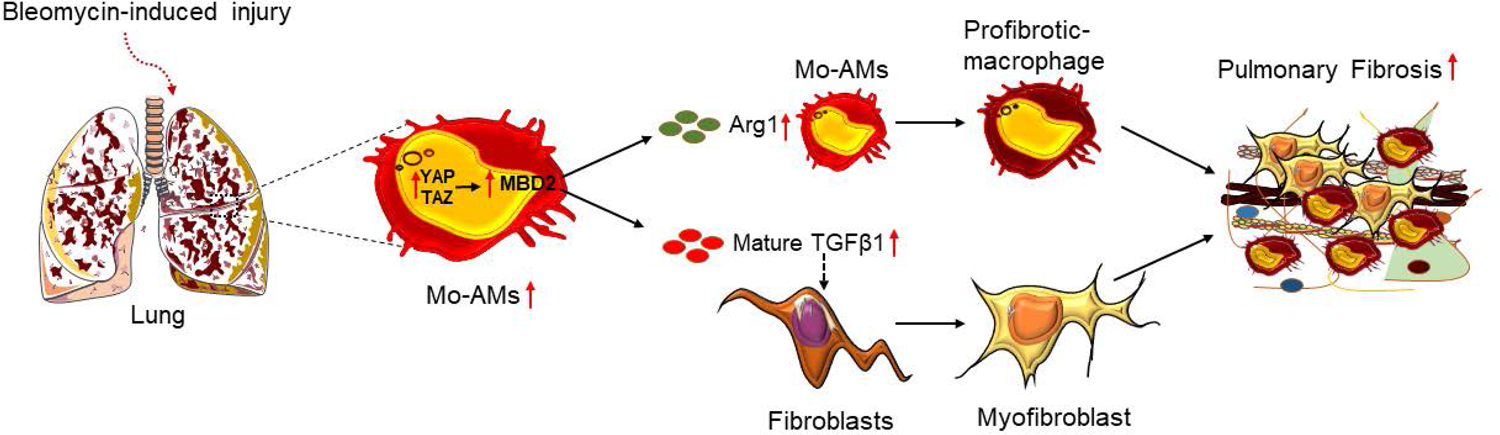
Graphical model for the macrophage –mediated role of YAP/TAZ during pulmonary fibrosis. Schematic representation showing that development of fibrosis associated with the recruitments of Mo-AMs after bleomycin treatment. PF augments the activation of YAP and TAZ in Mo-AMs which mediates the induction of MBD2 and ARG1 as well as release of pro-fibrotic cytokine TGFβ1 that activate macrophages and fibroblasts toward profibrotic phenotype leading to PF.

## Author Contributions

M.M.M., S.A.B.A.G., D.M.C., and H.B. performed experiments and analyzed data. W.S.F.W. provided reagents. M.M.M. designed the experiments, oversaw the entire project and wrote the manuscript. M.K.S. oversaw the entire project, designed the experiments, and edited the manuscript. All authors discussed the results and implications and commented on the manuscript at all stages.

## Acknowledgments

We are thankful to M.K.S. lab members for helpful discussion. This work was supported by funds from National Medical Research Council Singapore with grant number MOH-001130 (MOH-OFYIRG21nov-0005) to M.M.M and Duke-NUS Medical School Singapore and the Goh foundation and a Singapore National Research Foundation (NRF) fellowship (NRF-NRFF2016-01) to M.K.S.

---

**Supplementary Figure 1: Presence of macrophage increases during human lung fibrosis.** Immunofluorescence staining and quantification of CD68+ macrophages in collagen I+ area in lung sections from human patient diagnosed with pulmonary fibrosis compared with normal adult human lung (n = 3). The data are represented as the mean ± SEM; comparison by two-tailed unpaired t-test. *, *p* < 0.05; **, *p* < 0.01; ***, *p* < 0.001; NS, not significant.

**Supplementary Figure 2: Bleomycin treatment activates the YAP/TAZ expression in lung macrophages.** (A-B) Representative images from lung tissue sections after bleomycin-induced injury stained for CD68 (green), YAP (red), TAZ (pink), and Dapi (blue). Lung macrophages showed increased nuclear presence and activity of YAP and TAZ at 7 days after bleomycin treatment in control mice. In *dKO* lung, the expression of YAP and TAZ were inactivated in CD68^+^ macrophages. Green arrows indicate the cells that are positive for CD68 only; red arrows indicate the cells that are positive for YAP only; pink arrows indicate the cells that are positive for TAZ only; yellow arrows indicate the cells that are positive for CD68/YAP/Dapi (A) or CD68/TAZ/Dapi (B).

**Supplementary Figure 3: Working model showing bleomycin treatment induces lung fibrosis.** *Yap^flox/flox^*;*Taz^flox/flox^*, *LysM^cre^*;*Yap^flox/flox^*;*Taz^flox/flox^*, *R26^YAP5SA^* and *LysM^Cre^;R26^YAP5SA^* mice were exposed to intratracheal bleomycin administration and performed immunohistological analysis on lung sections. Histology of whole section of lungs from *Yap^flox/flox^;Taz^flox/flox^* mice at Day 7 and 14 post-fibrosis induced by bleomycin compared saline treated animals considered as Day 0. Lung tissues were assessed by Sirius Red and Trichrome staining to observe the fibrosis.

**Supplementary Figure 4: Loss of *Yap/Taz* in macrophage attenuates myofibroblasts and collagen production induced bleomycin.** (A-B) Immunofluorescence staining and quantification of αSMA and collagen I in lung sections from control and *dKO* mice (n = 5) treated with saline or bleomycin for 14 days. Fluorescence intensity was quantified with imageJ software. (C) Immunoblot analysis and quantification of αSMA and collagen I protein using lung tissue from control and *dKO* mice (n = 5) treated with saline or bleomycin for 14 days. The data are represented as the mean ± SEM; comparison by two-tailed unpaired t-test. *, *p* < 0.05; **, *p* < 0.01; ***, *p* < 0.001; NS, not significant.

**Supplementary Figure 5: Loss of *Yap/Taz* in macrophage improves alveolar epithelial repair after bleomycin injury.** (A) Representative plot from flow cytometry analysis to determine the changes in the percentage of AT2-cells identified as CD45^−^CD31^−^CD326^+^ MHCII^+^ in control *versus dKO* mice treated with bleomycin for 14 days. (B) mRNA levels AT2 cell marker *Sftpc* from control and *dKO* mice (n = 5-8) treated with saline or bleomycin for 14 days. (C) mRNA levels AT1 cell marker *Cav1* from control and *dKO* mice (n = 5-8) treated with saline or bleomycin for 14 days. The data are represented as the mean ± SEM; comparison by two-tailed unpaired t-test. *, *p* < 0.05; **, *p* < 0.01; ***, *p* < 0.001; NS, not significant.

**Supplementary Figure 6: YAP is hyperactivated in lung macrophages after bleomycin treatment in *LysM^Cre^;R26^YAP5SA^*mice.** Representative images and quantification from lung tissue sections after sham (saline) or bleomycin-induced injury stained for CD68 (green), YAP (red), and Dapi (blue). *LysM^Cre^;R26^YAP5SA^*lungs showed increased nuclear presence and activity of YAP at 14 days either with saline or bleomycin treatment compared to *R26^YAP5SA^* lungs; although the nuclear localization and activity of YAP was more prominent after bleomycin-induced injury. Green arrows indicate the cells that are positive for CD68 only; red arrows indicate the cells that are positive for YAP only; yellow arrows indicate the cells that are positive for CD68, YAP, and Dapi. The data are represented as the mean ± SEM; comparison by two-tailed unpaired t-test. *, *p* < 0.05; **, *p* < 0.01; ***, *p* < 0.001; NS, not significant.

**Supplementary Figure 7: Overexpression of *Yap* in macrophage impairs alveolar epithelial repair after bleomycin injury.** (A) mRNA levels AT2 cell marker *Sftpc* from *R26^YAP5SA^*and *LysM^Cre^;R26^YAP5SA^* mice (n = 3-4) treated with saline or bleomycin for 14 days. (B) mRNA levels AT1 cell marker *Cav1* from *R26^YAP5SA^* and *LysM^Cre^;R26^YAP5SA^* mice (n = 3-4) treated with saline or bleomycin for 14 days. The data are represented as the mean ± SEM; comparison by two-tailed unpaired t-test. *, *p* < 0.05; **, *p* < 0.01; ***, *p* < 0.001; NS, not significant.

**Supplementary Figure 8: Loss of *Yap/Taz* in macrophage ameliorates MBD2 expression after bleomycin injury.** (A) Immunofluorescence staining to show the co-localization of YAP or TAZ with MBD2 in CD68+ macrophages in lung sections from human patients diagnosed with pulmonary fibrosis. (B) Immunofluorescence staining to show the co-localization of YAP or TAZ with MBD2 in CD68+ macrophages in lung sections from bleomycin-induced mice. (C) Immunostaining and quantification of MBD2+ macrophages marked with CD68 in lung tissue sections from control and *dKO* mice (n = 5) treated with bleomycin for 14 days. (D) Immunostaining and quantification of MBD2+ macrophages marked with CD68 in lung tissue sections from *R26^YAP5SA^* and *LysM^Cre^;R26^YAP5SA^*mice (n = 5) treated with bleomycin for 14 days. The data are represented as the mean ± SEM; comparison by two-tailed unpaired t-test. *, *p* < 0.05; **, *p* < 0.01; ***, *p* < 0.001; NS, not significant.

## References

Adamson IY, Bowden DH (1974) The type 2 cell as progenitor of alveolar epithelial regeneration. A cytodynamic study in mice after exposure to oxygen. Lab Invest 30: 35–42

Ai K, Pan J, Zhang P, Li H, He Z, Zhang H, Li X, Li Y, Yi L, Kang Y, Wang Y, Xiang X, Chai X, Zhang D (2022) Methyl-CpG-binding domain protein 2 contributes to renal fibrosis through promoting polarized M1 macrophages. Cell Death Dis 13: 125

Antoniades HN, Neville-Golden J, Galanopoulos T, Kradin RL, Valente AJ, Graves DT (1992) Expression of monocyte chemoattractant protein 1 mRNA in human idiopathic pulmonary fibrosis. Proc Natl Acad Sci U S A 89: 5371–5

Aran D, Looney AP, Liu L, Wu E, Fong V, Hsu A, Chak S, Naikawadi RP, Wolters PJ, Abate AR, Butte AJ, Bhattacharya M (2019) Reference-based analysis of lung single-cell sequencing reveals a transitional profibrotic macrophage. Nat Immunol 20: 163–172

Barkauskas CE, Cronce MJ, Rackley CR, Bowie EJ, Keene DR, Stripp BR, Randell SH, Noble PW, Hogan BL (2013) Type 2 alveolar cells are stem cells in adult lung. J Clin Invest 123: 3025–36

Barratt SL, Creamer A, Hayton C, Chaudhuri N (2018) Idiopathic Pulmonary Fibrosis (IPF): An Overview. J Clin Med 7

Boutanquoi PM, Burgy O, Beltramo G, Bellaye PS, Dondaine L, Marcion G, Pommerolle L, Vadel A, Spanjaard M, Demidov O, Mailleux A, Crestani B, Kolb M, Garrido C, Goirand F, Bonniaud P (2020) TRIM33 prevents pulmonary fibrosis by impairing TGF-beta1 signalling. Eur Respir J 55

Byrne AJ, Maher TM, Lloyd CM (2016) Pulmonary Macrophages: A New Therapeutic Pathway in Fibrosing Lung Disease? Trends Mol Med 22: 303–316

Chakarov S, Lim HY, Tan L, Lim SY, See P, Lum J, Zhang XM, Foo S, Nakamizo S, Duan K, Kong WT, Gentek R, Balachander A, Carbajo D, Bleriot C, Malleret B, Tam JKC, Baig S, Shabeer M, Toh SES et al. (2019) Two distinct interstitial macrophage populations coexist across tissues in specific subtissular niches. Science 363

Chen JY, Qiao K, Liu F, Wu B, Xu X, Jiao GQ, Lu RG, Li HX, Zhao J, Huang J, Yang Y, Lu XJ, Li JS, Jiang SY, Wang DP, Hu CX, Wang GL, Huang DX, Jiao GH, Wei D et al. (2020) Lung transplantation as therapeutic option in acute respiratory distress syndrome for coronavirus disease 2019-related pulmonary fibrosis. Chin Med J (Engl) 133: 1390–1396

Cui H, Jiang D, Banerjee S, Xie N, Kulkarni T, Liu RM, Duncan SR, Liu G (2020) Monocyte-derived alveolar macrophage apolipoprotein E participates in pulmonary fibrosis resolution. JCI Insight 5

Fu V, Plouffe SW, Guan KL (2017) The Hippo pathway in organ development, homeostasis, and regeneration. Curr Opin Cell Biol 49: 99–107

Gharaee-Kermani M, McCullumsmith RE, Charo IF, Kunkel SL, Phan SH (2003) CC-chemokine receptor 2 required for bleomycin-induced pulmonary fibrosis. Cytokine 24: 266–76

Gibbings SL, Goyal R, Desch AN, Leach SM, Prabagar M, Atif SM, Bratton DL, Janssen W, Jakubzick CV (2015) Transcriptome analysis highlights the conserved difference between embryonic and postnatal-derived alveolar macrophages. Blood 126: 1357–66

Gibbons MA, MacKinnon AC, Ramachandran P, Dhaliwal K, Duffin R, Phythian-Adams AT, van Rooijen N, Haslett C, Howie SE, Simpson AJ, Hirani N, Gauldie J, Iredale JP, Sethi T, Forbes SJ (2011) Ly6Chi monocytes direct alternatively activated profibrotic macrophage regulation of lung fibrosis. Am J Respir Crit Care Med 184: 569–81

Guo X, Zhao Y, Yan H, Yang Y, Shen S, Dai X, Ji X, Ji F, Gong XG, Li L, Bai X, Feng XH, Liang T, Ji J, Chen L, Wang H, Zhao B (2017) Single tumor-initiating cells evade immune clearance by recruiting type II macrophages. Genes Dev 31: 247–259

Hagenbeek TJ, Webster JD, Kljavin NM, Chang MT, Pham T, Lee HJ, Klijn C, Cai AG, Totpal K, Ravishankar B, Yang N, Lee DH, Walsh KB, Hatzivassiliou G, de la Cruz CC, Gould SE, Wu X, Lee WP, Yang S, Zhang Z et al. (2018) The Hippo pathway effector TAZ induces TEAD-dependent liver inflammation and tumors. Sci Signal 11

Hou J, Shi J, Chen L, Lv Z, Chen X, Cao H, Xiang Z, Han X (2018) M2 macrophages promote myofibroblast differentiation of LR-MSCs and are associated with pulmonary fibrogenesis. Cell Commun Signal 16: 89

Hussell T, Bell TJ (2014) Alveolar macrophages: plasticity in a tissue-specific context. Nat Rev Immunol 14: 81–93

Idiopathic Pulmonary Fibrosis Clinical Research N, Raghu G, Anstrom KJ, King TE, Jr., Lasky JA, Martinez FJ (2012) Prednisone, azathioprine, and N-acetylcysteine for pulmonary fibrosis. N Engl J Med 366: 1968–77

Iyonaga K, Takeya M, Saita N, Sakamoto O, Yoshimura T, Ando M, Takahashi K (1994) Monocyte chemoattractant protein-1 in idiopathic pulmonary fibrosis and other interstitial lung diseases. Hum Pathol 25: 455–63

Jager B, Seeliger B, Terwolbeck O, Warnecke G, Welte T, Muller M, Bode C, Prasse A (2021) The NLRP3-Inflammasome-Caspase-1 Pathway Is Upregulated in Idiopathic Pulmonary Fibrosis and Acute Exacerbations and Is Inducible by Apoptotic A549 Cells. Front Immunol 12: 642855

Janssen WJ, Barthel L, Muldrow A, Oberley-Deegan RE, Kearns MT, Jakubzick C, Henson PM (2011) Fas determines differential fates of resident and recruited macrophages during resolution of acute lung injury. Am J Respir Crit Care Med 184: 547–60

Joshi N, Watanabe S, Verma R, Jablonski RP, Chen CI, Cheresh P, Markov NS, Reyfman PA, McQuattie-Pimentel AC, Sichizya L, Lu Z, Piseaux-Aillon R, Kirchenbuechler D, Flozak AS, Gottardi CJ, Cuda CM, Perlman H, Jain M, Kamp DW, Budinger GRS et al. (2020) A spatially restricted fibrotic niche in pulmonary fibrosis is sustained by M-CSF/M-CSFR signalling in monocyte-derived alveolar macrophages. Eur Respir J 55

Kayagaki N, Warming S, Lamkanfi M, Vande Walle L, Louie S, Dong J, Newton K, Qu Y, Liu J, Heldens S, Zhang J, Lee WP, Roose-Girma M, Dixit VM (2011) Non-canonical inflammasome activation targets caspase-11. Nature 479: 117–21

LaCanna R, Liccardo D, Zhang P, Tragesser L, Wang Y, Cao T, Chapman HA, Morrisey EE, Shen H, Koch WJ, Kosmider B, Wolfson MR, Tian Y (2019) Yap/Taz regulate alveolar regeneration and resolution of lung inflammation. J Clin Invest 129: 2107–2122

Lasithiotaki I, Giannarakis I, Tsitoura E, Samara KD, Margaritopoulos GA, Choulaki C, Vasarmidi E, Tzanakis N, Voloudaki A, Sidiropoulos P, Siafakas NM, Antoniou KM (2016) NLRP3 inflammasome expression in idiopathic pulmonary fibrosis and rheumatoid lung. Eur Respir J 47: 910–8

Lederer DJ, Martinez FJ (2018) Idiopathic Pulmonary Fibrosis. N Engl J Med 378: 1811–1823

Ley B, Collard HR, King TE, Jr. (2011) Clinical course and prediction of survival in idiopathic pulmonary fibrosis. Am J Respir Crit Care Med 183: 431–40

Lis-Lopez L, Bauset C, Seco-Cervera M, Cosin-Roger J (2021) Is the Macrophage Phenotype Determinant for Fibrosis Development? Biomedicines 9

Luppi F, Cerri S, Beghe B, Fabbri LM, Richeldi L (2004) Corticosteroid and immunomodulatory agents in idiopathic pulmonary fibrosis. Respir Med 98: 1035–44

McCubbrey AL, Barthel L, Mohning MP, Redente EF, Mould KJ, Thomas SM, Leach SM, Danhorn T, Gibbings SL, Jakubzick CV, Henson PM, Janssen WJ (2018) Deletion of c-FLIP from CD11b(hi) Macrophages Prevents Development of Bleomycin-induced Lung Fibrosis. Am J Respir Cell Mol Biol 58: 66–78

Mia MM, Cibi DM, Abdul Ghani SAB, Song W, Tee N, Ghosh S, Mao J, Olson EN, Singh MK (2020) YAP/TAZ deficiency reprograms macrophage phenotype and improves infarct healing and cardiac function after myocardial infarction. PLoS Biol 18: e3000941

Misharin AV, Morales-Nebreda L, Reyfman PA, Cuda CM, Walter JM, McQuattie-Pimentel AC, Chen CI, Anekalla KR, Joshi N, Williams KJN, Abdala-Valencia H, Yacoub TJ, Chi M, Chiu S, Gonzalez-Gonzalez FJ, Gates K, Lam AP, Nicholson TT, Homan PJ, Soberanes S et al. (2017) Monocyte-derived alveolar macrophages drive lung fibrosis and persist in the lung over the life span. J Exp Med 214: 2387–2404

Moore BB, Hogaboam CM (2008) Murine models of pulmonary fibrosis. Am J Physiol Lung Cell Mol Physiol 294: L152–60

Moore BB, Paine R, 3rd, Christensen PJ, Moore TA, Sitterding S, Ngan R, Wilke CA, Kuziel WA, Toews GB (2001) Protection from pulmonary fibrosis in the absence of CCR2 signaling. J Immunol 167: 4368–77

Noble PW, Albera C, Bradford WZ, Costabel U, Glassberg MK, Kardatzke D, King TE, Jr., Lancaster L, Sahn SA, Szwarcberg J, Valeyre D, du Bois RM, Group CS (2011) Pirfenidone in patients with idiopathic pulmonary fibrosis (CAPACITY): two randomised trials. Lancet 377: 1760–9

Okuma T, Terasaki Y, Kaikita K, Kobayashi H, Kuziel WA, Kawasuji M, Takeya M (2004) C-C chemokine receptor 2 (CCR2) deficiency improves bleomycin-induced pulmonary fibrosis by attenuation of both macrophage infiltration and production of macrophage-derived matrix metalloproteinases. J Pathol 204: 594–604

Osterholzer JJ, Olszewski MA, Murdock BJ, Chen GH, Erb-Downward JR, Subbotina N, Browning K, Lin Y, Morey RE, Dayrit JK, Horowitz JC, Simon RH, Sisson TH (2013) Implicating exudate macrophages and Ly-6C(high) monocytes in CCR2-dependent lung fibrosis following gene-targeted alveolar injury. J Immunol 190: 3447–57

Redente EF, Keith RC, Janssen W, Henson PM, Ortiz LA, Downey GP, Bratton DL, Riches DW (2014) Tumor necrosis factor-alpha accelerates the resolution of established pulmonary fibrosis in mice by targeting profibrotic lung macrophages. Am J Respir Cell Mol Biol 50: 825–37

Richeldi L, du Bois RM, Raghu G, Azuma A, Brown KK, Costabel U, Cottin V, Flaherty KR, Hansell DM, Inoue Y, Kim DS, Kolb M, Nicholson AG, Noble PW, Selman M, Taniguchi H, Brun M, Le Maulf F, Girard M, Stowasser S et al. (2014) Efficacy and safety of nintedanib in idiopathic pulmonary fibrosis. N Engl J Med 370: 2071–82

Rose CE, Jr., Sung SS, Fu SM (2003) Significant involvement of CCL2 (MCP-1) in inflammatory disorders of the lung. Microcirculation 10: 273–88

Schwartz DA, Helmers RA, Dayton CS, Merchant RK, Hunninghake GW (1991) Determinants of bronchoalveolar lavage cellularity in idiopathic pulmonary fibrosis. J Appl Physiol (1985) 71: 1688–93

Schyns J, Bureau F, Marichal T (2018) Lung Interstitial Macrophages: Past, Present, and Future. J Immunol Res 2018: 5160794

Sefik E, Qu R, Junqueira C, Kaffe E, Mirza H, Zhao J, Brewer JR, Han A, Steach HR, Israelow B, Blackburn HN, Velazquez SE, Chen YG, Halene S, Iwasaki A, Meffre E, Nussenzweig M, Lieberman J, Wilen CB, Kluger Y et al. (2022) Inflammasome activation in infected macrophages drives COVID-19 pathology. Nature 606: 585–593

Selman M, Pardo A (2020) The leading role of epithelial cells in the pathogenesis of idiopathic pulmonary fibrosis. Cell Signal 66: 109482

Singh A, Chakraborty S, Wong SW, Hefner NA, Stuart A, Qadir AS, Mukhopadhyay A, Bachmaier K, Shin JW, Rehman J, Malik AB (2022) Nanoparticle targeting of de novo profibrotic macrophages mitigates lung fibrosis. Proc Natl Acad Sci U S A 119: e2121098119

Sisson TH, Mendez M, Choi K, Subbotina N, Courey A, Cunningham A, Dave A, Engelhardt JF, Liu X, White ES, Thannickal VJ, Moore BB, Christensen PJ, Simon RH (2010) Targeted injury of type II alveolar epithelial cells induces pulmonary fibrosis. Am J Respir Crit Care Med 181: 254–63

Smith RE, Strieter RM, Phan SH, Lukacs NW, Huffnagle GB, Wilke CA, Burdick MD, Lincoln P, Evanoff H, Kunkel SL (1994) Production and function of murine macrophage inflammatory protein-1 alpha in bleomycin-induced lung injury. J Immunol 153: 4704–12

Suga M, Iyonaga K, Ichiyasu H, Saita N, Yamasaki H, Ando M (1999) Clinical significance of MCP-1 levels in BALF and serum in patients with interstitial lung diseases. Eur Respir J 14: 376–82

Wang D, Zhang Y, Xu X, Wu J, Peng Y, Li J, Luo R, Huang L, Liu L, Yu S, Zhang N, Lu B, Zhao K (2021a) YAP promotes the activation of NLRP3 inflammasome via blocking K27-linked polyubiquitination of NLRP3. Nat Commun 12: 2674

Wang G, Lu X, Dey P, Deng P, Wu CC, Jiang S, Fang Z, Zhao K, Konaparthi R, Hua S, Zhang J, Li-Ning-Tapia EM, Kapoor A, Wu CJ, Patel NB, Guo Z, Ramamoorthy V, Tieu TN, Heffernan T, Zhao D et al. (2016) Targeting YAP-Dependent MDSC Infiltration Impairs Tumor Progression. Cancer Discov 6: 80–95

Wang S, Xie F, Chu F, Zhang Z, Yang B, Dai T, Gao L, Wang L, Ling L, Jia J, van Dam H, Jin J, Zhang L, Zhou F (2017) YAP antagonizes innate antiviral immunity and is targeted for lysosomal degradation through IKKvarepsilon-mediated phosphorylation. Nat Immunol 18: 733–743

Wang Y, Zhang L, Huang T, Wu GR, Zhou Q, Wang FX, Chen LM, Sun F, Lv Y, Xiong F, Zhang S, Yu Q, Yang P, Gu W, Xu Y, Zhao J, Zhang H, Xiong W, Wang CY (2022) The methyl-CpG-binding domain 2 facilitates pulmonary fibrosis by orchestrating fibroblast to myofibroblast differentiation. Eur Respir J 60

Wang Y, Zhang L, Wu GR, Zhou Q, Yue H, Rao LZ, Yuan T, Mo B, Wang FX, Chen LM, Sun F, Song J, Xiong F, Zhang S, Yu Q, Yang P, Xu Y, Zhao J, Zhang H, Xiong W et al. (2021b) MBD2 serves as a viable target against pulmonary fibrosis by inhibiting macrophage M2 program. Sci Adv 7

Wood KH, Zhou Z (2016) Emerging Molecular and Biological Functions of MBD2, a Reader of DNA Methylation. Front Genet 7: 93

Wu J, Yan Z, Schwartz DE, Yu J, Malik AB, Hu G (2013) Activation of NLRP3 inflammasome in alveolar macrophages contributes to mechanical stretch-induced lung inflammation and injury. J Immunol 190: 3590–9

Wuyts WA, Agostini C, Antoniou KM, Bouros D, Chambers RC, Cottin V, Egan JJ, Lambrecht BN, Lories R, Parfrey H, Prasse A, Robalo-Cordeiro C, Verbeken E, Verschakelen JA, Wells AU, Verleden GM (2013) The pathogenesis of pulmonary fibrosis: a moving target. Eur Respir J 41: 1207–18

Zhou X, Li W, Wang S, Zhang P, Wang Q, Xiao J, Zhang C, Zheng X, Xu X, Xue S, Hui L, Ji H, Wei B, Wang H (2019) YAP Aggravates Inflammatory Bowel Disease by Regulating M1/M2 Macrophage Polarization and Gut Microbial Homeostasis. Cell Rep 27: 1176–1189 e5

